# Learning transcriptome architecture from sequence with a long-context RNA foundation model

**DOI:** 10.1101/2024.08.26.609813

**Authors:** Ali Saberi, Benedict Choi, Shaopu Zhou, Mohsen Naghipourfar, Simai Wang, Aldo Hernández-Corchado, Rohan Dandage, Najia Bouaddouch, Arsham Mikaeili Namini, Ali Poursina, Vijay Ramani, Amin Emad, Hamed S. Najafabadi, Hani Goodarzi

## Abstract

Linking DNA sequence to genomic function remains one of the grand challenges in genetics and genomics. Here, we present a large-scale compendium of single-molecule transcriptome sequencing of diverse cancer cell lines, revealing their isoform diversity and specificity. We used this compendium to build Mach-1, an RNA foundation model that learns how the nucleotide sequence of unspliced pre-mRNA dictates transcriptome architecture—the relative abundances and molecular structures of mRNA isoforms. By using the Striped-Hyena architecture, Mach-1 handles extremely long sequence inputs at nucleotide resolution (64 kilobase pairs), allowing for quantitative, zero-shot prediction of all aspects of transcriptome architecture, spanning isoform abundance, structure, and variant-induced splicing changes. To test both the interpretive and generative capabilities of Mach-1, we experimentally validated its learned regulatory grammar and predictions through perturbation of RNA-binding proteins nominated by Mach-1 to impact targeted splicing, precise CRISPR editing of variants of uncertain significance that the model predicted to alter splicing, and *de novo* transcript synthesis and expression in human cells. Together, this release establishes a new foundation for sequence-to-transcript modeling. Mach-1’s representations can be extended and fine-tuned across a spectrum of biological contexts, from variant interpretation to RNA engineering.

## 1. Introduction

Cell identity and behavior in higher eukaryotes is chiefly determined by transcriptome architecture—the quantitative levels (i.e., transcript abundances) and molecular structures (i.e., constitutively and alternatively spliced exons) of RNA isoforms (Nilsen and Graveley, 2010). This architecture is ultimately encoded by the genomic sequence, as noncoding cis-regulatory elements within and around genes specify transcription start and end sites, splicing decisions, and overall expression levels of RNA molecules (Tao et al., 2024). Disruptions in these regulatory programs are common in complex diseases such as cancer, underscoring the need for technologies that can learn how genomic sequence and other factors shape the transcriptome (Wang and Burge, 2008).

Recent advances in deep representation learning have significantly advanced computational biology, enabling models that capture complex molecular processes such as protein-DNA binding, chromatin accessibility, RNA splicing, RNA polyadenylation site selection, and post-transcriptional regulation. Large-scale pre-trained (“foundation”) models have further extended these capabilities, offering versatile tools to model data across various domains; in genomics specifically, foundation models like Evo 2 (Brixi et al., 2025), Nucleotide Transformer (Dalla-Torre et al., 2025) and DNABERT-2 (Zhou et al., 2023) have achieved strong zero-shot performance on DNA-based prediction tasks. However, a model that jointly represents the full genomic context and the corresponding RNA molecular structures, at single-nucleotide resolution, remains lacking.

To addresses this gap, two key components are needed: a comprehensive molecular resource capturing transcriptome architecture across diverse genomic sequences and cellular contexts, and a model architecture capable of capturing their length and complexity. Short-read RNA sequencing has long been the dominant source of data for inferring the transcriptome architecture, but its limited read length (∼10^2^ bp) obscures the connectivity between transcription start sites, spliced exons, and 3’ end sites. This connectivity is especially important in light of known correlations between *e.g.* transcription initiation sites, first exon usage (Fiszbein et al., 2019), and transcription termination sites (Calvo-Roitberg et al., 2025). To overcome these limitations, we generated a new long-read transcriptomic dataset from 26 cancer cell lines representing ten tissues, pairing high-quality long-read Iso-Seq data for isoform discovery with RNA-matched short-read RNA-seq for accurate quantification. We combine our dataset with additional human and mouse samples to assemble a compendium of roughly 100 million full-length reads encompassing over 7 billion training tokens, forming a strong foundation modeling transcriptome architecture directly from sequence.

Prior RNA modeling efforts have relied on two major architectures. Convolutional neural networks, including Borzoi (Linder et al., 2025) and SpliceAI (Jaganathan et al., 2019)), can model long genomic regions through residual dilated convolutions or attention layers, but they provide only coarse-grained representations of long-range dependencies and rely on short-read data for training. Transformer-based models have emerged in recent years as an alternative, using pre-training objectives such as masked language modeling (MLM) or causal language modeling (CLM). For example, models such as SpliceBERT (Chen et al., 2024) employ MLM to learn RNA grammar from gene annotations, but its quadratic attention scaling limits context length and prevents modeling of full-length pre-mRNA. Architectures that incorporate Hyena operators with CLM, such as HyenaDNA (Nguyen et al., 2023) and Evo 2 (Brixi et al., 2025), enable efficient processing of long sequences but remain genome-centric and are not designed to represent the transcriptome’s isoform-level architecture.

In this study, we introduce Mach-1, the first self-supervised RNA foundation model that learns and predicts transcriptome architecture directly from long-context sequences. Using causal language modeling and an expanded RNA token set, Mach-1 is trained on our large-scale full-length transcriptome dataset to capture the relationship between genomic sequence and isoform structure. The model quantitatively predicts transcriptome architecture, including transcript start and end sites, exon composition, and isoform abundance, and produces informative embeddings and likelihoods for individual isoforms, overcoming the limitations of DNA-focused genome models in representing RNA diversity. Mach-1 outperforms existing long-context genome models in several zero-shot prediction tasks: (1) absolute and relative isoform abundances for a given pre-mRNA, (2) *in silico* exon trapping predictions as a stringent out-of-distribution test, and (3) the impact of sequence variants on splicing. Mach-1’s isoform embeddings can also be used to train downstream models that predict isoform abundance and context-specific isoform expression, revealing that these representations encode *cis*-regulatory programs. Finally, as a generative model, Mach-1 can synthesize realistic transcript sequences whose coding and non-coding features closely mirror those of natural mRNAs.

Importantly, we performed a series of experimental validations to systematically test Mach-1’s predictions and its learned understanding of the splicing grammar. We first showed that isoform sequencing following knockdown of RNA-binding proteins nominated by Mach-1 produced splicing changes consistent with those predicted. We then used CRISPR-based genome editing to confirm the functional impact of single-nucleotide variants previously annotated as variants of unknown significance (VUS) but identified by Mach-1 as regulators of splicing. Finally, to assess its generative capabilities, we synthesized *de novo* transcripts designed by Mach-1 and confirmed their expression and predicted splicing in human cells. Taken together, these results establish Mach-1 as a long-context RNA foundation model that captures both the predictive and generative grammar underlying transcriptome architecture.

## 2. Results

### 2.1. Building a map of human cell line transcriptomes using long-read RNA sequencing

To obtain a comprehensive view of transcriptomic architecture across diverse cellular contexts, we generated a large Iso-Seq (isoform-sequencing) dataset using the PacBio long-read RNA sequencing platform (Au et al., 2013). Specifically, we profiled 26 human cancer cell lines derived from ten tissue types, including skin, lung, breast, liver, kidney, pancreas, colon, prostate, bone, and bone marrow (Figure 1A). We refer to these collective datasets as the **C**ancer **C**ell-**L**ine **L**ong-read (C2L2) compendium. C2L2 comprises 35 million long reads with a median read length of 3 kilobases and a maximum read length of 16 kilobases (Figure 1B), corresponding to an average of 1.5 million reads per sample (Figure 1C). In parallel, we performed RNA-matched short-read RNA sequencing for all samples, generating an average of 55 million reads per sample (Figure S1A). Together, these paired datasets enabled systematic annotation and quantification of transcripts at isoform resolution (see Methods, Figure S1B).

**Figure 1.**
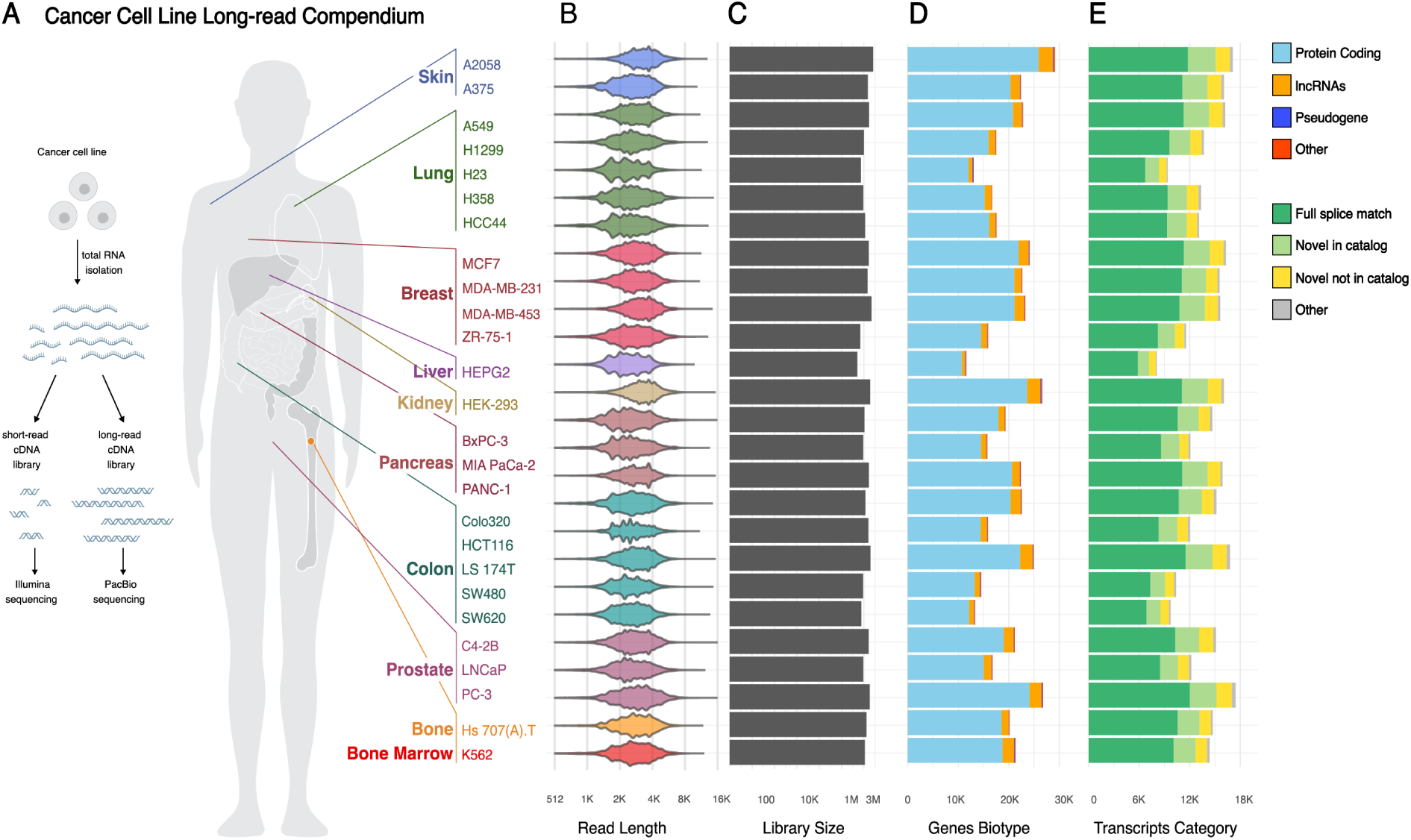
Overview of the Human Cancer Cell-Line Long-Read Compendium. **(A)** Experimental design and tissue origins of the Cancer Cell Line Long-read Compendium. (Left) Schematic showing the generation of matched short-read (Illumina) and long-read (PacBio) cDNA sequencing libraries from total RNA isolated from each cell line. (Right) Map of the 26 sampled cell lines, colored by their tissue of origin. **(B)** Violin plot of long-read lengths across cell lines. Each violin represents the distribution of read lengths for a given cell line, where the width reflects the density of observations at a given length. Wider regions indicate more frequent read lengths, while narrower regions indicate fewer observations. **(C)** Barplot of library sizes for each cell line, indicating the total number of reads obtained (log_10_ scale). **(D)** Stacked barplot of gene biotypes expressed (i.e., with at least one observed long read) in each cell line. **(E)** Stacked barplot categorizing inferred transcripts by structure, including full splice match, novel in catalog (novel combination of annotated splice junctions), and novel not in catalog (transcripts with at least one unannotated splice junction, following the notation used in (Pardo-Palacios et al., 2024).

Having established this comprehensive long-reads dataset, we next examined its overall composition and transcript diversity. As expected, the vast majority of detected transcripts were polyadenylated, with approximately 99% of inferred transcripts containing poly(A) tails. These include protein-coding genes, long non-coding RNAs (lncRNAs), and transcribed pseudogenes (Figure 1D). The majority of transcripts in the compendium correspond to previously annotated isoforms (Figure 1E). Nonetheless, we identified 1,307 novel genes alongside 23,691 known ones, and 10,139 novel transcripts in addition to 90,826 previously annotated isoforms. Most of these novel transcripts represent new combinations of known splice junctions, highlighting long-range interactions in RNA processing uncovered by Iso-Seq (Figure 1E).

### 2.2. C2L2 reveals patterns of cell-type specific isoform usage

Having established the C2L2 compendium, we next explored the transcriptomic diversity it captures across human cell lines. To characterize isoform expression patterns within the C2L2 dataset, we first assessed the global expression similarities among the 26 cell lines. Correlation analysis of log-transformed transcript per million (TPM) values demonstrated several key features of the data (Figure 2A). Biological replicates of the same cell line consistently show the highest correlation with each other, underscoring both the high reproducibility and cell line specificity of our measurements. Across the dataset, 22 of 26 cell lines showed a mean Pearson correlation greater than 0.7 between replicates (Figure S2A), and cell lines from the same tissue of origin frequently clustered together—for example, H23 and H1299 (lung) or SW480 and SW620 (colon)—further supporting the reproducibility and biological coherence of the dataset. Principal Component Analysis (PCA) revealed that transcriptomic profiles are structured by biological context, a pattern also captured by Uniform Manifold Approximation and Projection (UMAP) visualization (Figures S2B-E). Quantitative variance decomposition, performed by fitting a linear model to each principal component with metadata factors as covariates, showed that cell line identity and tissue of origin are the dominant drivers of this structure. Together, cell line identity and tissue of origin explain over 80% of the variance across the top principal components (Figure S2F), while technical batch effects contribute less than 5%.

**Figure 2.**
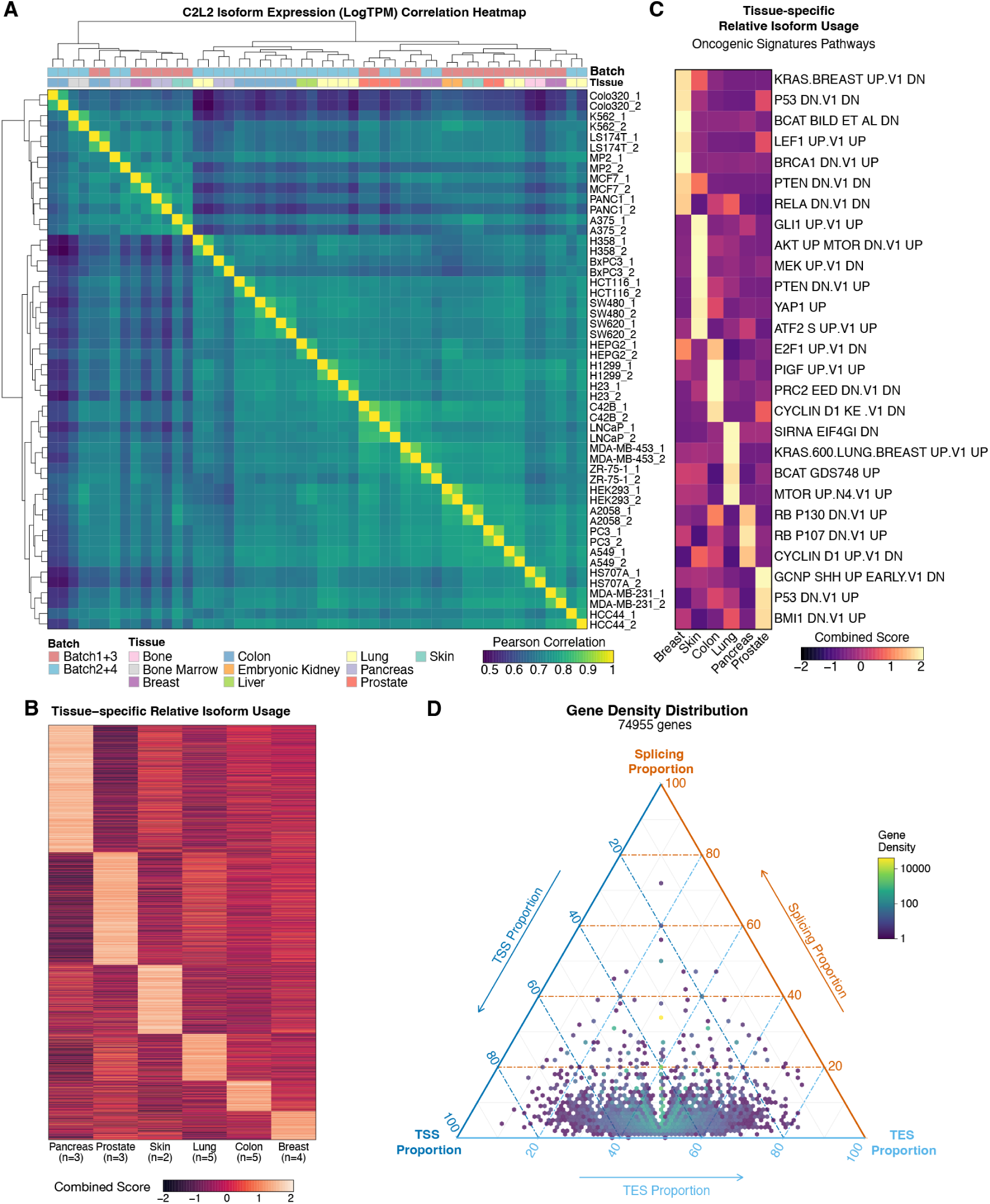
Tissue-specific isoform expression patterns and functional enrichment analysis. **(A)** Pearson correlation heatmap of log-transformed TPM expression values across 26 cell lines from the C2L2 dataset. Cell lines are clustered using Ward’s D2 hierarchical clustering. Annotation bars indicate tissue of origin (colored by tissue type) and experimental batch. **(B)** Heatmap of tissue-specific relative isoform usage, displaying beta regression coefficients for isoforms with significant differential expression (FDR < 0.1). Relative isoform usage is defined as the proportion of a transcript’s abundance relative to the total abundance of all isoforms from the same gene. Each row represents a transcript, and columns correspond to tissues. Transcripts are ordered diagonally —that is, sorted by enrichment scores so that tissue-specific transcripts align along the diagonal—to highlight tissue-specificity. **(C)** Functional enrichment heatmap for isoforms with tissue-specific relative usage against the MSigDB Oncogenic Signatures (C6) collection. The combined score, representing the product of −log_10_(p-value) and the z-score from EnrichR (Xie et al., 2021), is shown. Pathways are ordered diagonally to maximize the visibility of tissue-specific signals. **(D)** Ternary density plot illustrating the distribution of gene structure complexity for 74,955 genes in the inferred reference transcriptome. Each point represents a gene positioned according to the normalized contribution of these three mechanisms, with density shown on a log_10_ scale.

To elucidate the functional implications of these tissue-specific expression patterns, we first identified isoforms with significant tissue-specific *relative usage*—defined as the proportion of a transcript’s abundance relative to the total abundance of all isoforms from the same gene (Figure 2B). This metric captures isoform regulation independent of overall gene expression, providing a sensitive measure of post-transcriptional control that complements traditional differential expression analyses. We then performed pathway enrichment analysis, which revealed that tissue-specific isoform usage is functionally organized around key developmental and regulatory processes (Figure 2C; complete heatmap in Figure S2G). For example, PRC2 (SUZ12/EZH2)–mediated epigenetic repression was enriched in prostate cancer tissues, consistent with the established connection between PRC2 and disease progression (Heinke, 2023), suggesting that PRC2 may also undergo tissue-specific splicing regulation in prostate cancer. BMI1-driven self-renewal programs were likewise prominent in prostate models, reflecting BMI1’s role in neoplastic transformation (Yoo et al., 2016). In pancreas-derived lines, VEGF-mediated angiogenic signatures were notably enriched, aligning with the role of tumor vascularization in PDAC progression (Annese et al., 2019). In lung adenocarcinoma, KRAS-associated signaling pathways were enriched, matching KRAS mutation-linked expression profiles (Nagy et al., 2017). Together, these results indicate that isoform-level regulation reflects coherent, tissue-linked functional programs across diverse cellular contexts.

To provide a complementary perspective, we performed a parallel analysis based on absolute isoform abundance to identify tissue-specific transcripts. Unlike relative usage, which measures the proportion of each isoform within its parent gene, and therefore reflects post-transcriptional regulation, absolute abundance captures overall transcript levels also driven by transcriptional activity. A subsequent pathway enrichment yielded functional signatures that were concordant with those from the relative usage analysis (compare Figures S2G-H). This concordance is exemplified by the consistent identification of PRC2- and BMI1-related pathways in prostate models and KRAS-related signatures in lung cell lines across both methods. We then extended this dual analysis from the tissue level to individual cell lines. Functional enrichment on these cell-line-specific isoform sets again revealed coherent biological themes that remained consistent between the two metrics (compare Figures S2I-J). The recurrence of the same functional enrichments from two distinct metrics—relative usage, which primarily reflects post-transcriptional processes, and absolute abundance, which is more indicative of transcriptional activity—suggests a coordinated program of transcriptional and post-transcriptional regulation that reinforces cell identity. For example, pathways related to transcriptional regulation by TP53 are enriched at the tissue level in breast and colon, while MYC-related pathways are enriched at the cell-line level across both relative and absolute abundance analyses. The consistency of these biological signals across different analytical scales underscores the robustness of the information captured in the C2L2 dataset.

To systematically quantify the primary sources of transcript diversity, we used Cerberus (Reese, 2023) to perform a ternary simplex analysis that decomposes the complexity of each gene into three components: alternative transcription start sites (TSS), alternative transcription end sites (TES), and alternative splicing. This analysis, applied to 74,955 expressed genes defined based on combined evidence from both short- and long-read sequencing in our comprehensively annotated transcriptome, positioned each gene based on the normalized contributions of these three mechanisms to overall isoform diversity. The resulting density plot revealed the landscape of mechanisms through which genes generate transcript diversity (Figure 2D). To quantify these patterns, we classified each of the genes based on its dominant mechanism of diversity. The majority of genes (65,193; 87.0%) do not rely on a single dominant mechanism, instead generating diversity through a complex interplay of TSS, TES, and splicing variation, as indicated by their position away from the plot vertices. Among genes exhibiting a dominant mode of variation, diversity arising from alternative transcription start sites and alternative transcription end sites were similarly prevalent, accounting for 4,844 (6.5%) and 4,906 (6.5%) of genes, respectively. In contrast, only a small fraction of genes (12; 0.02%) relied primarily on alternative splicing as their main source of diversity. Overall, this distribution demonstrates that the combinatorial action of transcription initiation, polyadenylation site selection, and splicing is the predominant mechanism shaping isoform diversity across the human transcriptome—emphasizing the need for modeling frameworks that consider the full transcript structure rather than isolated splicing events.

### 2.3. Training a long-context RNA foundation model using causal language modeling

We designed Mach-1 as a transcriptomic model capable of handling full-length pre-mRNA sequences at nucleotide resolution, together with the molecular structure of their resulting mature mRNAs (Figure 3A). Mach-1 is based on the StripedHyena architecture (Poli et al., 2023b), which interleaves Hyena layers— data-controlled convolutional operators—with multi-head attention layers enhanced by rotary position embeddings. This hybrid design enables Mach-1 to process sequences up to 64K nucleotides in length, sufficient to capture most pre-mRNAs (191,273 of 263,073 isoforms in our compendium). Beyond its long-context capability, Mach-1 differs from previous StripedHyena-based genome foundation models (Nguyen et al., 2024) in being explicitly *splice-aware*. A key innovation in training Mach-1 is its tokenization scheme, which allows the model to represent RNA at single-nucleotide resolution while distinguishing between functional sequence regions (i.e. exonic, intronic, and untranscribed regions). Mach-1 employs a specialized vocabulary of 16 effective tokens: exonic regions are encoded in uppercase (A, C, U, and G), intronic regions in lowercase (a, c, u, and g), and non-RNA flanking sequences in special tokens (W, X, Y, and Z). Tokens H and M denote human and mouse species, respectively, while sentinel tokens S and P represent the transcription start site (TSS) and polyadenylation site (PAS). To standardize isoform lengths, Mach-1 adds upstream and downstream DNA sequences based on the longest isoform per gene, ensuring isoforms have uniform length and enabling direct comparison of their likelihoods (Figure 3B).

**Figure 3.**
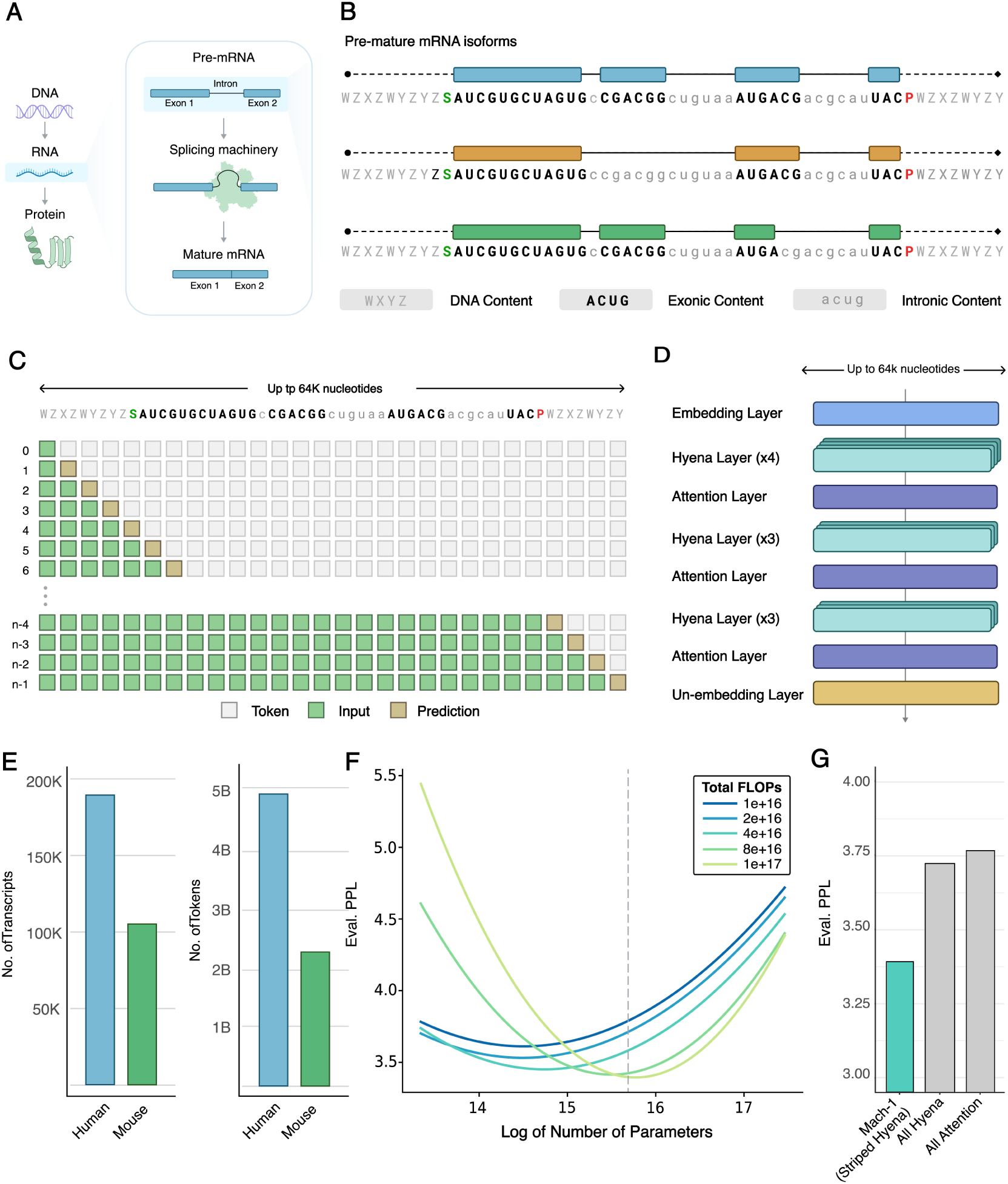
Mach-1 Model Architecture, Training, and Performance. **(A)** Schematic illustrating the transcription of pre-mRNA and the role of splicing machinery, which recognizes specific sequence features across the pre-mRNA to detect splice sites. The machinery then removes introns and joins exons to form mature mRNA. **(B)** Toy example demonstrating Mach-1’s tokenization approach given the genomic sequence and the exon-intron structure of different isoforms. **(C)** Overview of the causal language modeling used to train Mach-1, integrating Hyena layers with attention mechanisms to accommodate long nucleotide sequences. **(D)** Summary of training data, displaying number of transcripts and tokens for human and mouse datasets. **(E)** Scaling law analysis showing how evaluation perplexity decreases with increasing model size, measured in a natural log number of parameters (ranging from 0.5-40 million). Curves correspond to different computational budgets (1×10^16^–1×10^17^ FLOPs), illustrating the trade-off between model size and training cost. Beyond an optimal point, additional scaling leads to overfitting and reduced performance. **(F)** Bar plot comparing model performance for StripedHyena, Hyena-only, and attention-only architectures. Lower perplexity indicates better performance. **(G)** Modular design of the Mach-1 architecture, combining embedding, Hyena, and attention layers to efficiently process sequences up to 64K nucleotides efficiently.

Mach-1 was trained using causal language modeling (CLM) with a next-token prediction (NTP) objective (Figure 3C). This approach enables the model to learn the sequential dependencies inherent in RNA sequences by predicting each nucleotide based on its preceding context, a critical feature for tasks like isoform abundance prediction and variant effect analysis.

For pre-training, we compiled a large dataset from long-read RNA-seq data across human cancer cell lines. This included 52 samples (2 replicates for 26 cell lines) from the C2L2 compendium. Additionally, we incorporated 149 human and mouse samples from other long-read studies, spanning normal tissues, different cell types, COVID-19 studies, control conditions, and drug-treated models. In total, the dataset comprised 201 human samples, yielding approximately 85 million Full-Length Non-Chimeric (FLNC) reads and 277,481 unique transcripts, of which 191,273 had more than one exon and a standardized maximum length of 64K nucleotides. To augment cross-species learning, we added 46 mouse samples, contributing approximately 9.5 million FLNC reads and 153,133 unique transcripts, of which 106,453 had more than one exon and a maximum length of 64K nucleotides (Figure S3A-D). Altogether, Mach-1 was trained on 297,726 unique transcripts, totaling more than 7 billion training tokens (Figure 3D).

We next conducted a systematic scaling law analysis to determine the optimal balance among model size, training data, and computational resources. Consistent with prior studies, performance improved with increasing model size up to a limit, after which additional scaling yielded diminishing returns (Figure 3E). We also performed ablation studies by benchmarking Mach-1 against attention-only and Hyena-only architectures. Mach-1 consistently outperformed both alternatives, demonstrating the advantage of combining Hyena layers with attention mechanisms to process long RNA sequences while maintaining fine-grained contextual understanding (Figure 3F, Figure S3E). Based on these analyses, we selected a model with 16 blocks, width of 256 dimensions, and 6.5 million parameters. The architecture includes 13 Hyena layers interspersed with 3 rotary self-attention layers, providing an optimal trade-off between efficiency and predictive accuracy (Figure 3G; see Methods).

### 2.4. Mach-1 learns splice sites and *cis*-regulatory sequence elements

To further evaluate Mach-1’s capabilities, we examined its ability to identify splice sites and recognize regulatory sequence features that influence exon inclusion. As expected, isoform likelihoods predicted by Mach-1 were highly sensitive to splicing signals, particularly sequences surrounding splice junctions. To probe this further, we analyzed an experimentally validated exon in the mouse *Grin1* gene, which is well known to be regulated by two downstream Rbfox1 binding sites. Mach-1 assigned significantly higher likelihoods to splice site signals and Rbfox1 binding sites in transcripts that included the exon compared to those that excluded it, indicating that the model recognizes both canonical splice signals and associated regulatory elements (Figure 4A, Figure S4A).

**Figure 4.**
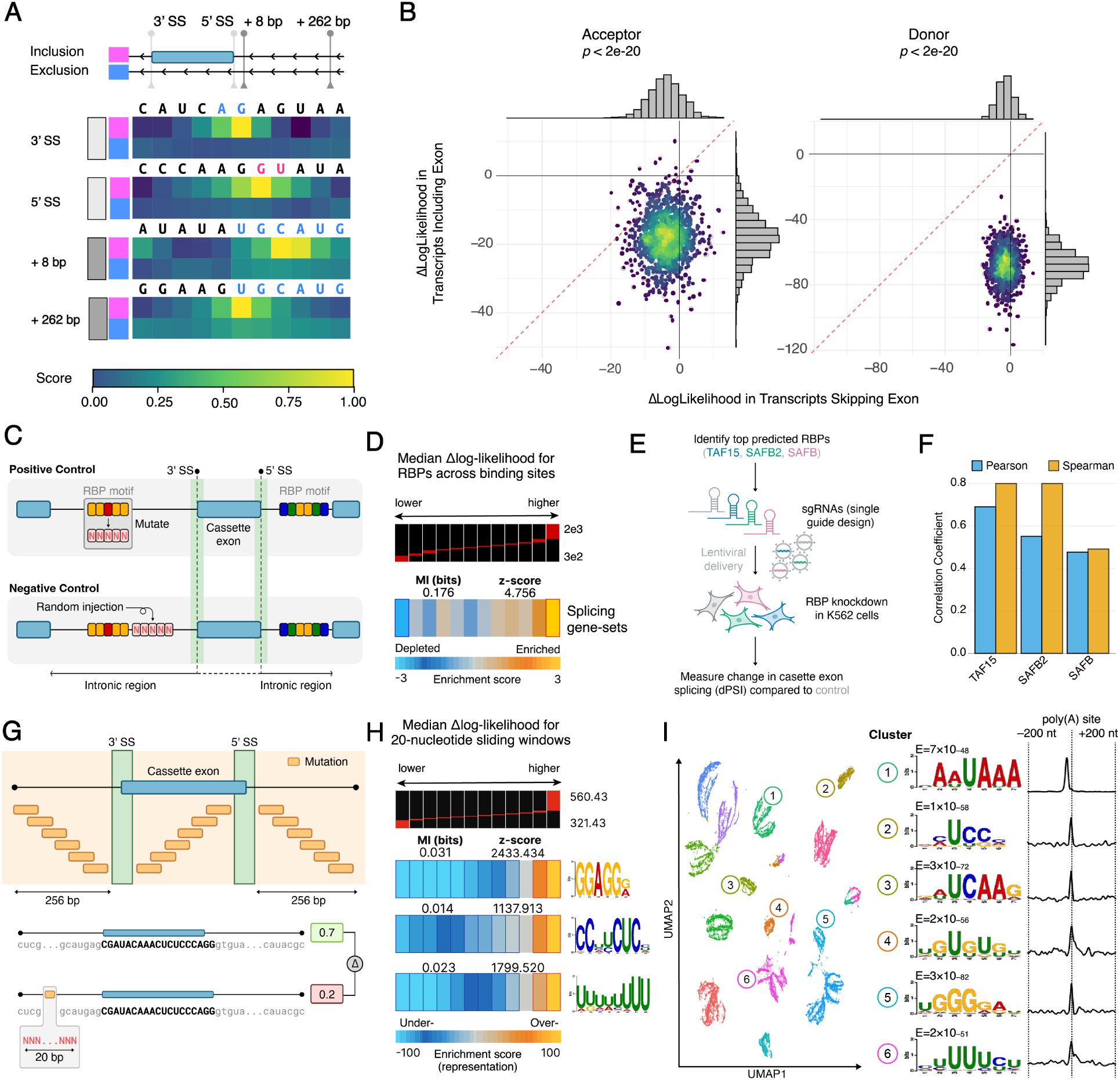
Mach-1 Learns Splice Sites and Cis-Regulatory Sequence Elements through Likelihood Estimation and Embedding Analyses. **(A)** Heatmaps showing average likelihoods for splice site signals and the Rbfox1 motif (+8 and +262 nucleotides upstream) matches in the *Grin1* gene, comparing transcripts with exon inclusion versus exclusion. In this experiment, each nucleotide in the sequence was systematically mutated into all possible alternatives, generating seven mutated sequences plus the original reference sequence. Mach-1 estimated the likelihood of each sequence, which was then normalized into a categorical probability distribution using the softmax function. The color scale highlights the probability of each sequence, where the only variation is a single nucleotide. A higher probability assigned to the sequence with the original reference nucleotide indicates that this nucleotide is critical for the model’s predictions, demonstrating the model’s sensitivity to specific nucleotide positions in RNA sequences. **(B)** Scatter plot illustrating the Δlog-likelihood (ΔLL) for the deletion of donor (right) and acceptor (left) splice site signals in isoforms that include a cassette exon (x-axis) versus those that exclude the exon (y-axis). Attached to each scatter plot is a histogram representation of the ΔLL distribution for each axis. The color scale of the scatter plot points represents density, with red indicating high density and blue indicating low density. Introns’ ΔLL were calculated using a one-sided Wilcoxon signed-rank test. **(C)** A schematic representation illustrating the methodology used to evaluate the impact of RNA-binding protein (RBP) motifs on splicing regulation. The positive set (top panel) shows the identification of RBP binding sites that overlap cassette exons and their flanking introns (excluding splice sites and ±5 nucleotides). Each binding site was then mutated (replaced with “N”), and the corresponding change in log-likelihood (ΔLL) of the isoform sequence was calculated. The negative set (bottom panel) involved selecting random regions of similar length that did not overlap with known RBP binding sites, which were also deleted to calculate ΔLL. The comparison between RBP-associated ΔLL and random sequence effects. **(D)** Enrichment and depletion of splicing factors, as a gene set, across the range of median Δlog-likelihood (ΔLL) across all RBPs. The heatmap displays the enrichment score (measured by log-transformed hypergeometric p-values [−log_1_0 p]) for splicing-related RBPs as a function of increasing ΔLL. The ΔLL values were divided into 12 equally populated bins, with each heatmap. **(E)** Schematic of the experimental validation of RBP-mediated splicing regulation. Following knockdown of a target RBP (TAF15, SAFB, or SAFB2) in K562 cells, changes in cassette exon inclusion (ΔPSI) were measured experimentally via short- and long-read RNA sequencing. These values were then correlated with Mach-1’s predicted change in log-likelihood (ΔLL) derived from the in silico mutation of the corresponding RBP’s binding sites in the flanking introns. **(F)** Bar plot showing the Pearson and Spearman correlation coefficients between Mach-1’s predicted ΔLL scores and the experimentally measured ΔPSI values for cassette exon events significantly altered by the knockdown of TAF15, SAFB, and SAFB2 (SUPPA2, FDR < 0.05). **(G)** Top: A schematic representation illustrating the sliding window approach used on a cassette exon and its flanking 256-bp intronic regions, excluding splice sites and ±5 nucleotides. A 20-bp window slides with a stride of 10 nucleotides over these regions, and for each window, the content is replaced with ‘N’. The Δlog-likelihood between the mutant (altered) and reference (unaltered) isoform sequences is calculated. Bottom: Among the eight significantly enriched motifs discovered, three were significantly matched (*q*-value < 0.05) to known splicing factors. The sequence logos represent these motifs and their associated splicing factors: SRSF1, PCBP2, and a combination of U2AF2 and TIA1. **(H)** Motif Enrichment Analysis Based on Δlog-likelihood Scores. Heatmaps display the enrichment (representation) of motifs from sliding windows with significant Δlog-likelihood decreases, with scores shown on a blue (underrepresented) to gold (overrepresented) color scale. Motif discovery identified three motifs matching known splicing factor binding preferences—SRSF1, PCBP2, and U2AF2/TIA1—accompanied by sequence logos. Mutual information (MI) values and Z-scores quantify the strength of association between motif presence and Δlog-likelihood changes, with Z-scores indicating how far the observed MI deviates from random expectations. The enrichment analysis is described in D. **(I)** UMAP visualization of PAS embeddings. For a few example clusters, the top enriched motifs near the poly(A) sites are shown, along with their positional biases. The e-values (E) indicate the statistical significance of motif enrichment, where lower values suggest stronger enrichment. The e-value, as reported by MEME, estimates the expected number of motifs with the same or higher log likelihood ratio, width, and site count in a similarly sized set of random sequences. Random sequences assume independent positions with letters chosen according to background frequencies.

To systematically assess Mach-1’s understanding of splicing code, we extracted all alternatively spliced cassette exon events from the C2L2-derived transcriptome. For each event, we selected two major isoforms: (1) the highest-expression isoform that included the exon and (2) the highest-expression isoform that excluded it. We then ablated the canonical two-nucleotide donor (GU) and acceptor (AG) splice site signals by replacing them with N and computed the resulting change in log-likelihood (Δlog-likelihood) relative to the reference isoform. As expected, Δlog-likelihood decreased significantly more (p < 1×10^−20^; one-sided Wilcoxon signed-rank test) for isoforms that included the cassette exon than those that excluded it, confirming that Mach-1 captures the sequence features distinguishing exon–intron boundaries and accurately models splicing outcomes (Figure 4B).

Next, we investigated whether Mach-1 also learns the influence of RNA-binding proteins (RBPs) on splicing.

Using ENCODE RBP binding data from K562 and HepG2 cells, we examined binding sites that overlapped cassette exons or their flanking introns, excluding regions within five nucleotides of splice donor or acceptor sites to remove the direct splice-site signal from the analysis and focus on additional regulatory influences. To simulate RBP binding loss, we replaced each binding site sequence with N and calculated Δlog-likelihood values relative to unaltered isoforms. As a negative control, we repeated the analysis using random regions of similar length that did not overlap known RBP binding sites (Figure 4C; Figure S4B). Sites with high Δlog-likelihood were significantly enriched for binding by known splicing regulators, suggesting that Mach-1 has internalized the functional impact of RBP-mediated splicing control (Figure 4D).

To directly test these predictions, we experimentally perturbed three RBPs, TAF15, SAFB, and SAFB2, that Mach-1 nominated as key regulators of specific splicing events. Using CRISPR interference (CRISPRi, Gilbert et al. (2014)), we knocked down each RBP in K562 cells and quantified the resulting splicing changes via short- and long-read RNA sequencing (Figure 4E). We reasoned that if Mach-1’s predictions are biologically meaningful, the change in log-likelihood (ΔLL) upon binding site ablation should correlate with the experimentally observed change in exon inclusion (ΔPSI) upon RBP depletion. As expected, for cassette exons significantly affected by RBP knockdown (FDR < 0.05), the change in exon inclusion (ΔPSI) correlated strongly with Mach-1’s predicted change in likelihood (ΔLL), demonstrating that the model’s predictions reflect causal regulatory effects (Figure 4F; Figure S4C). Correlations between ΔLL and transcript-level log-fold changes further supported this relationship (Figure S4D).

Because ENCODE covers only a subset of RBPs (Van Nostrand et al., 2016), we next sought to identify additional *cis*-regulatory elements underlying Mach-1’s predictions. For this, we applied a sliding-window ablation analysis (a 20-nt window, 10-nt stride) across cassette exons and their 256-bp flanking introns, excluding ±5 nucleotides around splice sites. For each window, we replaced the sequence with Ns and calculated the resulting Δlog-likelihood between the ablated and reference sequences (Figure 4G). Windows showing the largest decreases in log-likelihood (top 50%) were analyzed with the MEME suite for *de novo* motif discovery (Machanick and Bailey, 2011). Eight significantly enriched motifs were identified and mapped to known RBPs using Tomtom (Gupta et al., 2007), including binding motifs for SRSF1, PCBP2, and a U-rich element consistent with U2AF2/TIA1 recognition. The strong enrichment of these motifs in high-impact regions confirms that Mach-1 learns the sequence grammar that governs splicing regulation (Figure 4H).

Finally, we examined whether Mach-1’s transcript-level embeddings capture additional layers of post-transcriptional regulation. Using the polyadenylation site (PAS) token embedding—treated as a sentinel token summarizing transcript context—we observed distinct clusters of transcript representations that correlated with regulatory motif composition. Some clusters were enriched for canonical polyadenylation signals (e.g., upstream AAUAAA motifs with downstream U- or GU-rich elements), while others contained less characterized motifs such as CUCC and AUCAA near the PAS (Figure 4I). These results demonstrate that Mach-1 captures both the predictive and mechanistic grammar underlying RNA processing, from splicing control to polyadenylation site selection.

### 2.5. Mach-1 generalizes in zero-shot mode across diverse transcriptomic tasks

Zero-shot prediction refers to a model’s ability to make accurate predictions on tasks it was never explicitly trained for. Such performance indicates the model has internalized the functional grammar of RNA sequences and can generalize to unseen predictive settings. To evaluate Mach-1’s generalization capacity, we tested it on several distinct downstream tasks encompassing major axes of transcriptome regulation: absolute isoform abundance prediction (or relative usage prediction), *in silico* exon trapping prediction, and splicing effect prediction for non-coding variants.

#### 2.5.1. Modeling isoform abundance from sequence using PAS likelihoods

Transcript abundance is determined by a combination of *cis*-regulatory features embedded in the genomic sequence that act at both the DNA and RNA levels. We evaluated Mach-1’s ability to predict mean isoform abundance across diverse cell lines using the PAS token likelihoods, which encapsulates the model’s understanding of the entire transcript. Isoforms with higher likelihoods should correspond to sequences and structural configurations more consistent with processed and expressed mRNAs; such isoforms are expected to be processed more frequently, which in turn results in higher mean abundance.

We computed the correlation between Mach-1’s predicted likelihoods and the observed relative isoform usage across genes. Mach-1 achieved a median of 0.22, higher than most state-of-the-art models, capturing the relationship between sequence composition and isoform output (Figure 5A). The only exception was Evo 2 (Brixi et al., 2025), which contains roughly 1000-fold more parameters than Mach-1. Notably, these results were obtained without any task-specific fine-tuning. A simple downstream linear model trained on PAS embeddings as isoform representations further increased the correlation with absolute mRNA abundance to 0.38 (Figure S5A), highlighting the interpretability and transferability of Mach-1’s learned representations.

**Figure 5.**
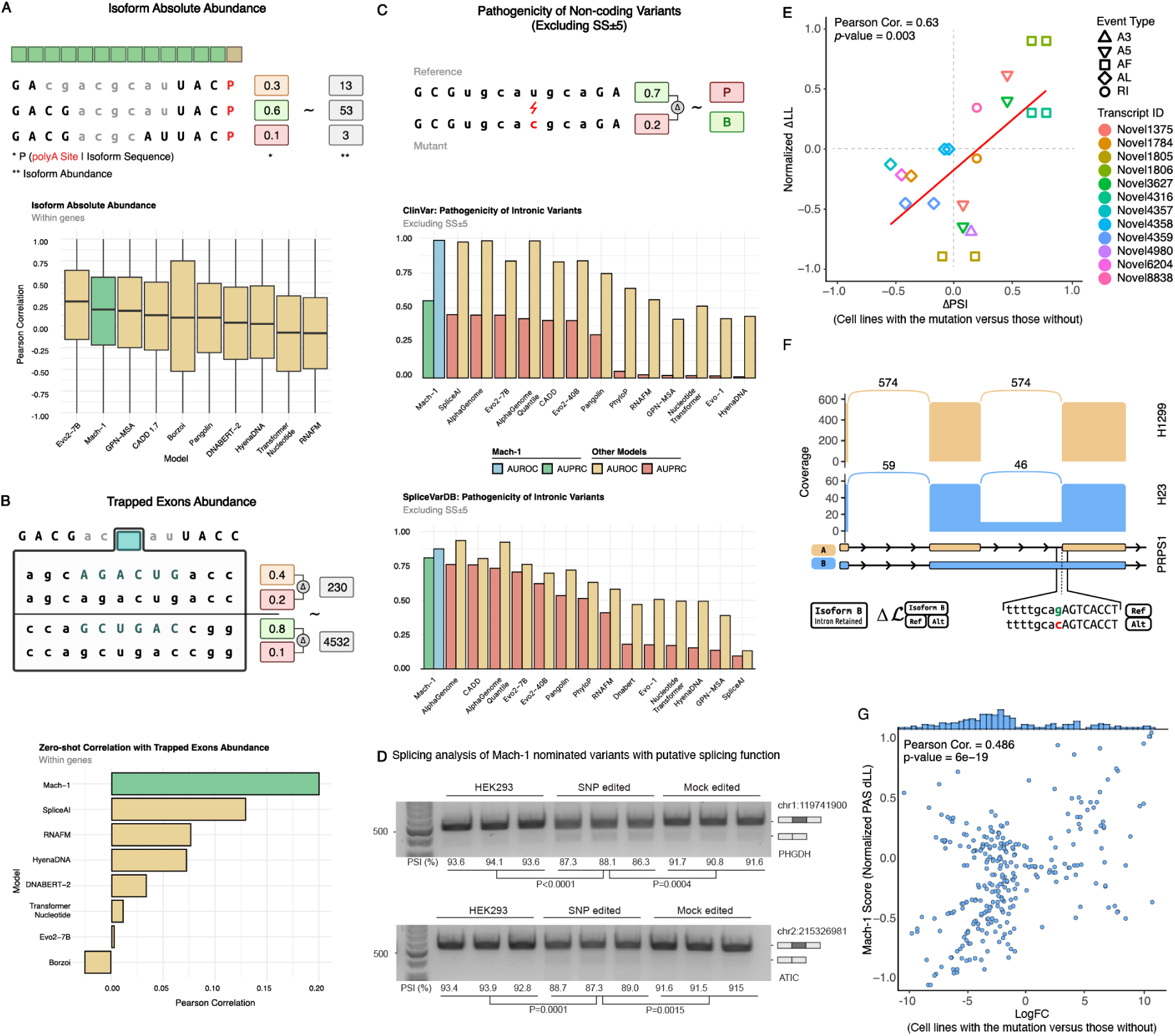
Model’s Zero-Shot and Trained Models Prediction Performance on Downstream Tasks. **(A)** Top: Schematic of the experiment for predicting isoform absolute abundance based on the PAS token likelihood, illustrating how the model estimates the probability of isoform sequences. Bottom: The boxplots show the correlation of the relative isoform usage versus likelihood within each gene. **(B)** Top: Schematic of the experiment for predicting trapped exon absolute abundance, detailing how the model evaluates exon inclusion. Bottom: The correlation between the Δlog-likelihood and the absolute abundance of trapped exons. **(C)** Top: Schematic of the experiment for predicting variant pathogenicity based on the difference in likelihoods between wild-type and mutant sequences. Bottom: The bar plots show the AUROC (left) and AUPRC (right). In each panel, Mach-1 is compared to other models on the same task. **(D)** Splicing analysis of Mach-1 nominated variants with putative splicing function. Alternative splicing patterns in CRISPR-edited cell lines were compared with the parental line via RT-PCR and gel electrophoresis. Distinct splicing isoforms showing a clear shift in splicing upon introduction of the variant. **(E)** Correlation between changes in normalized isoform likelihood (ΔLL) and changes in exon inclusion (ΔPSI) for novel alternative splicing events caused by somatic mutations in various cell lines. Each point represents a novel transcript with a distinct alternative splicing event type (A3, A5, AF, AL, or RI). **(F)** Comparison of isoform expression in H23 and H1299 lung cancer cell lines based on long-reads coverage for PRPS1. The sashimi plot (top) illustrates coverage differences between two PRPS1 isoforms in H23 (blue) and H1299 (yellow). The mutation that increases the relative likelihood of the isoform with the retained intron, based on Mach-1 predictions is shown at the bottom. **(G)** Correlation between the change in log-likelihood (ΔLL) predicted by Mach-1 and the log-fold change (logFC) in transcript expression for somatic mutations present in a subset of C2L2 cell lines. The ΔLL was calculated between the mutant and reference versions of the affected transcripts, while the logFC was calculated for the expression of these transcripts in cell lines with the mutation versus those without.

#### 2.5.2. Generalization to out-of-distribution splicing contexts via in silico exon trapping

A recent genome-wide exon-trapping screen identified approximately 1.25 million genomic fragments across the human genome capable of acting as autonomous exons, demonstrating that many sequences can be spliced when expressed in a permissive pre-mRNA context (Stepankiw et al., 2023). Because these fragments are drawn from unexpressed regions of the genome, this dataset serves as a stringent out-of-distribution (OOD) test of Mach-1’s ability to generalize beyond the sequences observed during training. We simulated the exon-trapping assay *in silico* by inserting each reported trapped exon, together with 500 nucleotides of flanking context, into the original experimental vector. Mach-1 then calculated likelihoods for the exon-included and exon-skipped configurations, and the resulting Δlog-likelihood values were compared to experimentally reported exon-trapping frequencies. The predicted and observed values were significantly correlated (Pearson *r* = 0.20, *p* = 2.1 × 10^−28^; Figure S5B), surpassing SpliceAI and other specialized baselines (Figure 5B). This strong performance on a truly OOD dataset demonstrates that Mach-1 has learned general, context-independent rules of splicing rather than memorizing transcriptomic patterns.

#### 2.5.3. Predicting functional impacts of non-coding variants without task-specific training

Non-coding genetic variants can perturb RNA processing and expression through regulatory mechanisms that do not alter protein sequence. There are many non-coding variants that fall within intronic sequences or untranslated regions of the gene—if functional, these variants can influence RNA processing and splicing, which may lead to pathogenic phenotypes. Accurate prediction of their functional consequences is critical for understanding disease risk, interpreting genome-wide association results, and improving clinical variant annotation.

We evaluated Mach-1’s zero-shot ability to distinguish pathogenic from benign or uncertain non-coding variants using high-confidence subsets from ClinVar (Landrum et al., 2014) (2+ star-rated variants) and SpliceVarDB (Sullivan et al., 2024), excluding canonical splice sites and their ±5-nt flanking regions. Pathogenicity was quantified as the likelihood difference between reference and mutant sequences. Mach-1 achieved AUROC/AUPRC values of 0.98/0.55 on ClinVar and 0.87/0.81 on SpliceVarDB, outperforming all other tested models (Figure 5C). These results, achieved using next-token prediction alone, highlight Mach-1’s internal representation of splicing-regulatory logic.

To experimentally validate these predictions, we selected six ClinVar variants of uncertain significance (VUS) that Mach-1 predicted to alter splicing and that fall close to annotated cassette exons. Each variant was precisely introduced into HEK293 cells using CRISPR–Cas9 (Figure S5C), and successful editing at the target loci was confirmed via Nanopore sequencing of genomic DNA (Figure S5D). We then assessed the impact of each edit on splicing by performing RT–PCR on RNA from the edited cell lines and comparing the resulting isoform patterns with those of both the parental (reference) and mock-edited control lines carrying identical editing scars. In two of six cases, the single-nucleotide substitution induced a clear and significant shift in the splicing pattern, resulting in exclusion of the proximal exon (Figure 5D). Notably, both of these experimentally validated splice-altering variants were classified as variants of uncertain significance in ClinVar and predicted to be benign by SpliceAI, highlighting Mach-1’s ability to uncover functional non-coding variants missed by existing prediction tools. One validated variant at the chr1:119741900 locus occurs at a site that is not evolutionarily conserved across vertebrates (Figure S5D), demonstrating that Mach-1 can identify functional regulatory variants even in regions lacking evidence of evolutionary constraint.

#### 2.5.4. Inferring cis-regulatory effects of somatic mutations on isoform expression

To assess Mach-1’s ability to predict how somatic mutations reshape transcriptome architecture, particularly through the induction of new alternative splicing events and transcripts isoforms, we leveraged cell line-specific somatic mutation data from the DepMap database (Arafeh et al., 2025) within the C2L2 cell line panel. Our goal was to determine whether Mach-1 could infer, in a zero-shot setting, how these mutations influence the likelihood of specific isoforms in each cell line without additional model training.

Using SUPPA2 (Trincado et al., 2018), we identified differentially spliced exons between pairs of cell lines within each tissue type. Focusing on mutations occurring near these differentially spliced events, we observed that the mutation-induced change in isoform likelihood closely tracked the experimentally observed change in isoform abundance. Across all events, the predicted and observed changes were significantly correlated (Pearson *r* = 0.63, *p* = 0.003; Figure 5E). As an illustrative example, a splice-acceptor mutation in *PRPS1* in the H23 lung cell line was predicted by Mach-1 to promote intron retention, a prediction confirmed by long-read RNA sequencing. The intron-retained isoform appeared exclusively in H23 cells but not in H1299 cells, consistent with Mach-1’s higher predicted likelihood for this transcript (Figure 5F).

We next extended this analysis to a broader set of somatic mutations present in a subset of C2L2 cell lines but absent in others of the same tissue lineage. For each affected transcript, we compared the change in isoform expression (log_2_ fold-change between mutant and wild-type lines) with the corresponding Mach-1–predicted change in log-likelihood (ΔLL). This analysis revealed a robust correlation (Pearson *r* = 0.48, *p* = 6 × 10^−19^; Figure 5G) between the model’s zero-shot predictions and measured expression shifts. Notably, in most cases, Mach-1 correctly predicted both the magnitude and direction of transcript abundance changes. However, a minority of variants showed strong predicted effects with incorrect directionality, indicating that while Mach-1 successfully identifies functionally impactful variants, incorporating contextual features such as chromatin state or trans-factor activity could further improve prediction of effect polarity.

Together, these results demonstrate that Mach-1 can infer the *cis*-regulatory consequences of somatic mutations directly from sequence, extending its zero-shot predictive capacity from generic splicing effects to quantitative modeling of mutation-driven transcriptome remodeling.

### 2.6. Leveraging Mach-1 embeddings to predict cell line–specific transcriptomes

Mach-1 was trained on human and mouse transcriptomes derived from diverse long-read RNA-seq datasets encompassing multiple cell lines and tissues. To test whether its learned representations capture biologically meaningful information beyond sequence-level prediction, we built cell line–specific models using Mach-1 embeddings as input features. This analysis allowed us to evaluate how well Mach-1’s internal representations encode regulatory information that enables accurate, context-specific predictions of isoform expression across distinct cellular environments.

Our hypothesis, consistent with our earlier observations, was that a collection of splicing factors govern RNA processing in each cell line, and that Mach-1 has learned the underlying *cis*-regulatory elements that allow accurate prediction of isoform likelihoods. To demonstrate that this regulatory grammar is captured by the model and linked to the *trans* factors active in every cell line, we used the PAS token embeddings of isoforms to train downstream cell line-specific models. Specifically, for each cell line, we trained a gradient boosting regressor to predict isoform abundance in that cell line based on its PAS token embeddings. The Pearson correlation coefficient between the observed abundance (logTPM) and the predicted abundance across 26 cell lines is shown in Figure 6A, ranging from 0.29 for H358 to 0.46 for PANC1 (Figure 6B), all statistically significant (maximum adjusted *p*-value = 1.4e–03). A comprehensive heatmap detailing the performance across all benchmarked models is provided in Figure S6A.

**Figure 6.**
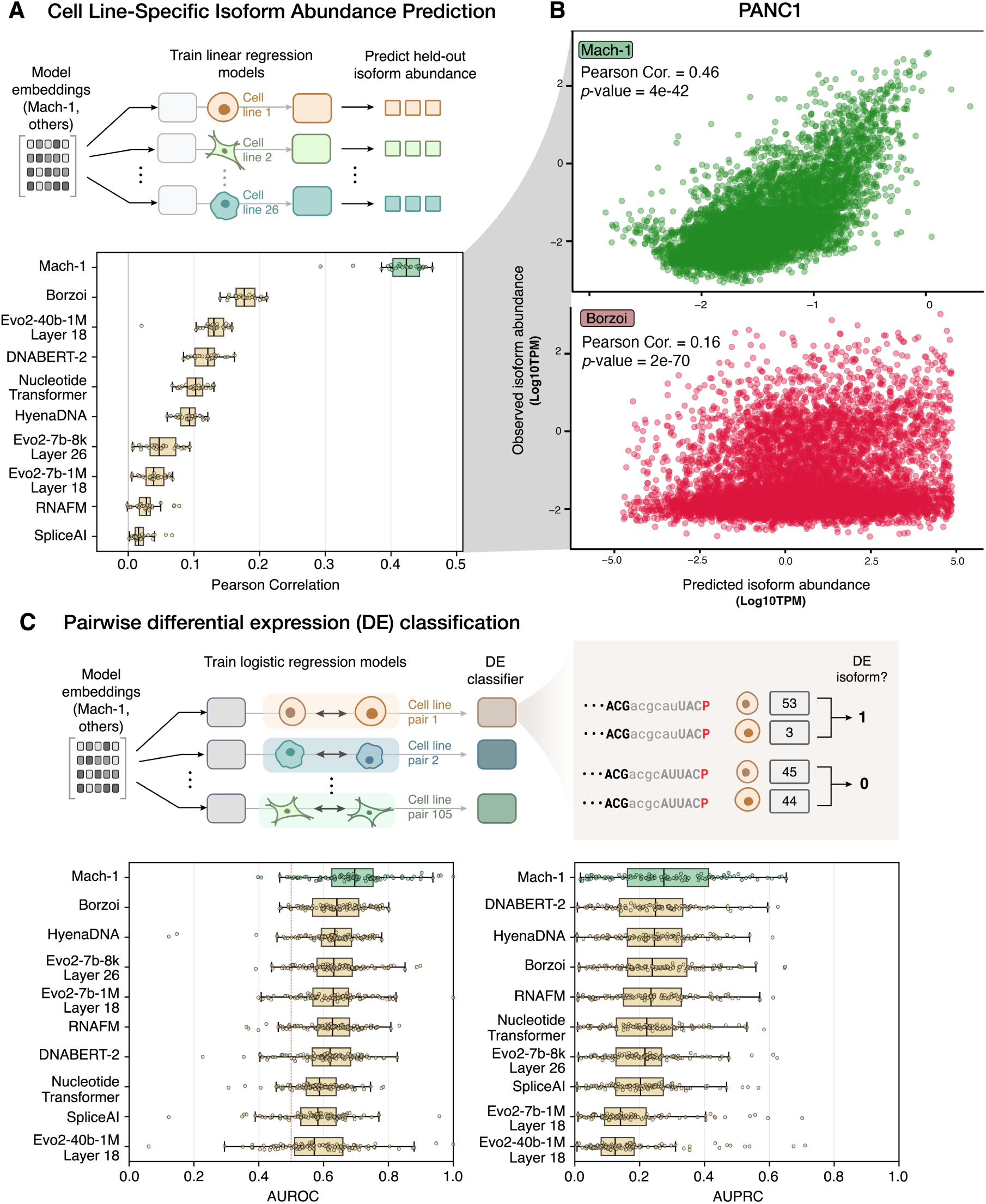
Mach-1 fine-tuned prediction performance on cell line–specific downstream tasks. **(A)** Pearson correlation coefficients between observed and predicted isoform abundance across 26 cell lines, each modeled using a gradient boosting regressor trained on PAS token embeddings. Correlation coefficients were calculated using isoforms encoded on chromosome 8, which was held out as a test set during model training and validation. Cell lines are colored by tissue of origin, representing ten tissue types in total. **(B)** Scatter plot of predicted versus observed isoform abundance for the PANC1 cell line test set, comparing Mach-1 (best model) with Borzoi (next-best model). Point density is indicated by color intensity, with darker regions representing higher density. **(C)** Boxplots showing the AUROC (left) and AUPRC (right) scores for prediction of differential isoform expression on the held-out chromosome 8 test set. Each box represents results from 105 pairwise comparisons between cell lines within the same tissue of origin. The distributions summarize performance across all comparisons.

To further evaluate Mach-1’s ability to capture *trans*-acting effects, we performed pairwise differential expression analysis between cell lines within the same tissue type (considering tissues represented by multiple lines). For each of the 105 cell line pairs, we trained a classifier using PAS token embeddings to predict whether a transcript was significantly upregulated in one cell line relative to the other (ground truth determined using LIMMA, with FDR < 0.05 and log_2_ fold-change > 1). Distributions of AUROC and AUPRC scores across all pairwise classifications are shown in Figure 6C. Mach-1 consistently outperformed alternative foundation models (Figure S6B), with detailed performance metrics (AUROC and AUPRC) provided in Figure S6C and S6D. Together, these analyses demonstrate that Mach-1’s embeddings capture the underlying *cis*- and *trans*-regulatory programs governing isoform expression, enabling accurate prediction of cell line–specific transcriptomic landscapes from sequence-derived representations alone.

### 2.7. Mach-1 generates synthetic transcripts that recapitulate human transcriptome architecture

To evaluate Mach-1’s ability to reproduce the structural complexity of the transcriptome, we prompted the model to generate approximately 50,000 transcripts. Generation was conditioned on the “HS” token, directing Mach-1 to synthesize human transcripts from the transcription start site (TSS) to the polyadenylation site (PAS), denoted by “S” and “P” tokens, respectively (Figure 7A). An equivalent number of annotated human transcripts were sampled as a reference dataset for comparison.

**Figure 7.**
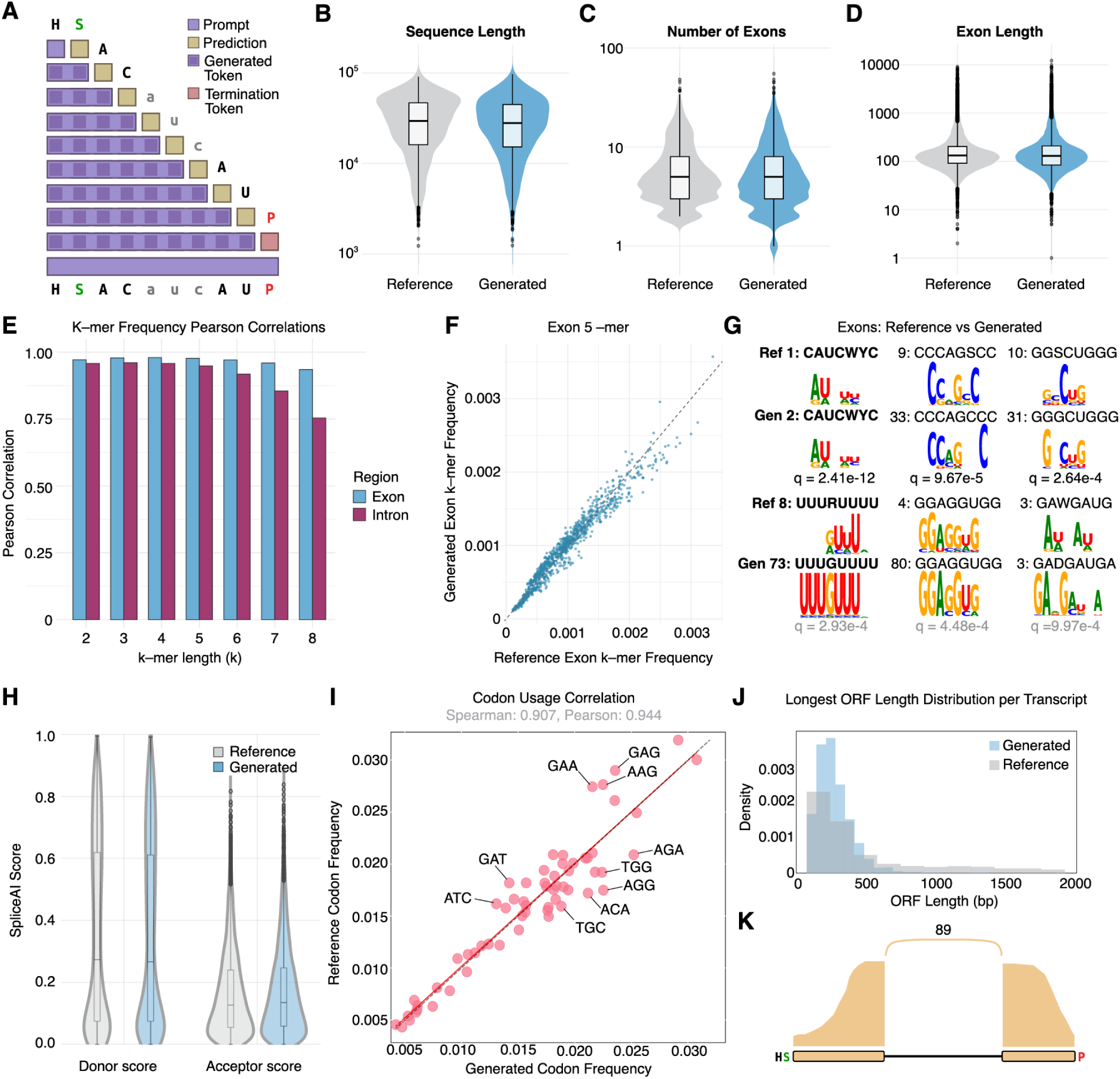
Mach-1 generates synthetic transcripts that recapitulate the architecture of the human transcriptome. **(A)** Schematic of the autoregressive process used by Mach-1 to generate transcript sequences. **(B-D)** Violin plots comparing the distributions of sequence length (B), number of exons (C), and exon length (D) between the generated (blue) and reference (gray) transcript sets. **(E)** Pearson correlation of k-mer frequencies (k=2 to 8) between generated and reference sequences, shown separately for exonic (purple) and intronic (teal) regions. **(F)** Scatter plot of 5-mer frequencies in the exonic regions of generated versus reference transcripts. **(G)** Sequence logos for the top three most enriched motifs identified in the exonic regions of reference transcripts (top row) and their matched counterparts in the generated transcripts (bottom row). **(H)** Violin plots comparing SpliceAI-predicted donor and acceptor site scores between generated and reference transcripts. **(I)** Scatter plot showing the correlation of codon usage frequencies between open reading frames (ORFs) from generated transcripts and human reference coding sequences. **(J)** Density plot of ORF lengths for generated and reference transcripts. **(K)** Sashimi plot from experimental validation showing a representative synthetic transcript expressed and spliced in HEK293 cells, consistent with the *in silico* design.

Our analysis revealed a high degree of structural similarity between the Mach-1-generated and reference transcriptomes. The length distributions of the full-length synthetic transcripts closely mirrored those of natural transcripts (Figure 7B). Likewise, Mach-1 accurately recapitulated the characteristic length distributions of exons and introns (Figure 7C, 7D, Figure S7A, S7B) and the overall exon-intron organization, demonstrating that the model learns to alternate effectively between functional regions. At the sequence level, we observed a high correlation in k-mer frequencies between reference and generated transcripts (Figure 7E, 7F, Figure S7C, S7D). Motif enrichment analysis further showed that over 85% of the motifs enriched in the reference sequences (MEME, E-value < 1e−5) had significant matches among motifs enriched in the Mach-1–generated sequences (Tomtom, *q*-value < 0.01) (Figure 7G, Figure S7E, S7F). Together, these findings indicate that Mach-1 not only reproduces the global composition of human transcripts but also captures recurrent regulatory motifs that shape transcriptome architecture.

To assess the functional viability of Mach-1–generated sequences, we performed a series of *in silico* analyses. Splice junctions in synthetic transcripts were largely indistinguishable from those in the reference set based on SpliceAI scores (Figure 7H), indicating that the generated transcripts are competent for correct splicing. Following *in silico* splicing to obtain mature mRNA sequences, we identified open reading frames (ORFs) and analyzed their codon usage. Codon frequencies in generated ORFs were highly correlated with those in reference human coding sequences (Spearman’s r = 0.907, Pearson’s r = 0.944; Figure 7I), showing that Mach-1 has learned the statistical structure of coding regions despite the relative scarcity of exonic tokens in the training data. Although the generated ORFs were, on average, shorter than reference ORFs (Figure 7J), these results were achieved entirely in a zero-shot setting, without hyperparameter tuning to optimize sequence length.

Finally, to provide experimental validation, we synthesized ten Mach-1–generated transcripts and introduced them into HEK293 cells. Subsequent sequencing confirmed that these synthetic transcripts were expressed and spliced with high fidelity to the model’s predicted exon–intron boundaries. In one representative construct, 89 junction-spanning reads supported the exact predicted splice junction, with no evidence of alternative boundaries (Figure 7K). Together, these findings demonstrate that Mach-1 can generate biologically coherent and functionally viable transcripts, establishing a foundation for future fine-tuning and applications in generative transcriptome design.

## 3. Discussion

Here, we present C2L2 and Mach-1, a first-in-class long-read transcriptomic resource and associated foundation model that predicts transcriptome architecture from input genomic sequence. The C2L2 long-read dataset—spanning 26 cancer cell lines across ten tissues—provides a comprehensive view of complete transcript structure and expression levels including the coordinated usage of transcription start sites, splicing patterns, and polyadenylation sites on individual molecules. By revealing intrinsic properties of the transcriptome that are inaccessible to short-read sequencing (e.g. observation of coordinated pairing between transcript 5^′^ and 3^′^ ends), C2L2 enables training of foundation models that learn the sequence basis for novel biological insights, such as the recent discovery that transcription initiation and termination are molecularly coupled processes (Calvo-Roitberg et al., 2025).

Mach-1, in turn, is such a foundation model. It leverages C2L2 to generate predictions of spliced mRNA sequences from up to ∼64,000 input DNA tokens. Importantly, Mach-1’s novelty is not simply owed to the unique training dataset it is built on, as it improves in two additional ways over existing state-of-the-art predictive models. First, Mach-1’s training paradigm and architecture are distinct from existing transformer-based approaches for predicting RNA splicing, such as SpliceAI and Borzoi, in that its model uses a splice-aware tokenization scheme and StripedHyena operators to model full-length pre-mRNA sequences. Second, this improvement offers Mach-1 distinct advantages over existing models, including high performance on zero-shot tasks such as the prediction of isoform abundance and structure, the inclusion of trapped exons in arbitrary sequence contexts, and functional assessment of non-coding variants acting over very long distances of pre-mRNA sequence. These advantages highlight a key difference between Mach-1 and other existing models: while models such as SpliceAI and Borzoi excel at predicting splicing patterns on the bases of RNA-seq coverage and splice site usage (tasks well suited to short-read sequencing data), Mach-1 is a causal language model, and learns probabilities of isoforms conditioned on long pre-mRNA sequences. This key difference drives Mach-1’s performance in predicting isoform patterns that are out-of-distribution with respect to training data.

We validate Mach-1’s knowledge of the rules of RNA splicing with extensive biological experimentation. Using CRISPR-based editing of Mach-1-predicted splicing variants, we achieve a 2/6 success rate of predicting splice-altering genetic variants that, importantly, were predicted to have no effect by SpliceAI. Notably, one of these functional variants is not evolutionarily conserved, underscoring Mach-1’s ability to detect regulatory signals beyond conservation-based heuristics. While a 33% hit rate is substantial given that these variants were tested outside of their native cellular contexts, it likely represents a conservative lower bound on Mach-1’s predictive power, as many sequence perturbations may require specific cell-type or trans-factor environments to manifest their effects. We provide some evidence that Mach-1 has learned key trans regulators of isoforms as well. By performing RNA binding protein (RBP) knockdown experiments motivated by motifs specifically prioritized by Mach-1 in isoform generation tasks, we find that experimentally measured splicing changes following CRISPRi-mediated knockdown match those predicted by Mach-1 following mutation of corresponding RBP motifs. Although the current model is cell-type agnostic, its learned representations can already be extended: we used Mach-1 embeddings to train cell line–specific predictors of isoform abundance and differential expression, revealing that these embeddings encode both general transcriptomic information and cell type–specific nuances.

While Mach-1 demonstrates strong predictive performance, our analysis of somatic mutation effects also highlights a common limitation at the frontier of sequence-based modeling. For a subset of variants, Mach-1 correctly captured the magnitude but not the direction of expression changes, a phenomenon also observed in other large-scale variant effect models (Huang et al., 2023; Sasse et al., 2023). This sign-agnostic behavior suggests that although current models can recognize the presence of regulatory perturbations, they remain limited in resolving context-dependent outcomes such as activation versus repression. Future iterations of Mach-1 could address this by integrating additional regulatory information, including cell state embeddings and trans-factor abundance, thereby capturing the full cellular context that modulates the directionality of transcriptomic responses.

We close by highlighting the unique framework for mechanistic interpretation offered by Mach-1. Unlike existing transformer-based architectures, Mach-1’s token-level likelihoods and learned embeddings are inherently suited for interpretable representations of regulatory grammar such as motifs, pairwise dependencies, and higher-exposing motifs, dependencies, and higher-order relationships among *cis*-regulatory elements. Such interpretability transforms Mach-1 from a purely predictive model into a hypothesis-generating platform capable of uncovering novel regulators of RNA processing and transcriptome structure. Looking ahead, we anticipate that Mach-1’s foundation-model architecture will also be well-suited for transfer learning. Because its embeddings capture the complexity and long-range sequence context of RNA processing, they can serve as general-purpose representations for downstream fine-tuning on diverse biological tasks (e.g. tissue-specific splicing prediction; disease-associated splice variant prediction; optimization of RNA-based therapeutics). Integration of Mach-1 with multimodal foundation models spanning chromatin accessibility, RNA structure probing, and transcriptional dynamics will ultimately offer a unified framework to model information flow across biology’s central dogma.

## 4. Methods

### 4.1. Data Collection

#### 4.1.1. Cell culture

All cells were cultured in a 37^◦^C 5% CO_2_ humidified incubator and in growth medium supplemented with antibiotics including penicillin (100 units/mL), streptomycin (100 *μ*g/mL) and amphotericin B (1 *μ*g/mL). The following cell lines were cultured in DMEM high-glucose medium (Gibco) supplemented with 10% fetal bovine serum (FBS), glucose (4.5 g/L), L-glutamine (4 mM) and sodium pyruvate (1 mM): MCF7 (ATCC HTB-22), MDA-MB-231 (ATCC HTB-26), MDA-MB-453 (ATCC HTB-131), LS174T (ATCC CL-188), A375 (CRL-1619), PANC-1 (ATCC CRL-1469), C4-2B (ATCC CRL-3315), Hs 707(A).T (ATCC CRL-7448), HEK-293 (ATCC CRL-1573) and A2058 (ATCC CRL-3601). The next cell lines were cultured in RPMI-1640 medium (Gibco) supplemented with 10% FBS, L-glutamine (2 mM) and HEPES (25 mM): BxPC-3 (ATCC CRL-1687), LNCaP (ATCC CRL-1740), ZR-75-1 (ATCC CRL-1500), H358 (ATCC CRL-5807), H1299 (ATCC CRL-5803), Colo320 (DZMC ACC-144), H23 (ATCC CRL-5800), HCC44 (DZMC ACC-534), MIA PaCa-2 (ATCC CRL-1420) and K562 (ATCC CCL-243). The cell lines including HCT116 (ATCC CCL-247), SW480 (ATCC CCL-228) and SW620 (ATCC CCL-227) were cultured in McCoy’s 5A (modified) medium (Gibco) with 10% FBS, glucose (3 g/L) and L-glutamine (1.5 mM). The lines A549 (ATCC CCL-185) and PC-3 (CRL-1435) were cultured in Ham’s F-12K medium (Gibco) supplemented with 10% FBS and L-glutamine (2 mM). The HEPG2 line (ATCC HB-8065) was cultured in Eagle’s Minimum Essential Medium (ATCC) supplemented with 10% FBS, L-glutamine (2 mM) and sodium pyruvate (1 mM). All cell lines were routinely screened for mycoplasma with a PCR-based assay and tested negative. They were also routinely authenticated using STR profiling at UC Berkeley Sequencing Facility.

#### 4.1.2. RNA isolation

Cells were collected and lysed in TRIzol reagent (ThermoFisher). The lysate was mixed with chloroform and phase-separated in a phaselock gel tube heavy (Quantabio) by centrifugation. The aqueous phase containing RNA was then mixed with one volume of 100% ethanol, and column-purified using Zymo QuickRNA Miniprep kit with in-column DNase treatment per the manufacturer’s protocol. RNA integrity (RIN ≥ 9) was confirmed using the 4200 Tapestation System (Agilent) prior to library construction.

#### 4.1.3. Short-read and long-read RNA-seq

Short-read RNA-seq libraries were prepared using the SMARTer Stranded Total RNA-seq kit v3 (TaKaRa Bio) following the manufacturer’s instructions. The libraries were sequenced on a NovaSeq 6000 instrument at UCSF Center for Advanced Technology. Long-read RNA-seq libraries were constructed using the NEBNext Single Cell/Low Input cDNA Synthesis & Amplification Module, PacBio Iso-Seq Express Oligo Kit and SMRTbell express template prep kit 3.0 according to the manufacturer’s protocol. The libraries were sequenced on a PacBio Revio at UCSF.

### 4.2. Data Preprocessing

We used an in-house pipeline for long-read RNA-seq data processing, incorporating several critical tools within *Nextflow* to ensure reproducible data preparation. The workflow began with the IsoSeq pipeline, starting with *pbccs*, which was used to generate HiFi reads using raw subreads through circular consensus sequencing (CCS). These high-quality reads were then processed using *lima* to demultiplex the sequences based on barcodes and to remove cDNA primers. The *isoseq-refine* tool was subsequently applied for further refinement, including polyA tail trimming and concatemer removal, resulting in Full-Length Non-Chimeric (FLNC) reads that capture complete transcript structures. Following the generation of FLNC reads, the data were aligned to the reference genome using *minimap2* (Li, 2018), employing the splice preset optimized for long-read spliced alignment. After alignment, *samtools* (Li et al., 2009) was used to sort and index the BAM files. For transcript isoform quantification, we utilized *bambu* (Chen et al., 2023), a tool specifically designed for long-read RNA-seq data, operating in both discovery and quantification modes. In the discovery mode, *bambu* performed error correction on the reads and extended existing annotations to identify novel transcripts, effectively inferring the transcriptome by discovering previously unannotated isoforms. In the subsequent quantification mode, *bambu* quantified transcript abundance by associating read classes with specific transcripts, including both known and novel isoforms. By orchestrating these tools within *Nextflow*, we were able to automate and streamline the entire data processing workflow to ensure greater consistency and reproducibility

#### 4.2.1. Correlation Analysis and Hierarchical Clustering

The correlation heatmap in Figure 2A was generated by computing Pearson correlation coefficients on log2-transformed TPM expression values across all 26 cell lines in the C2L2 dataset. Hierarchical clustering of both rows and columns was performed using the Ward’s D2 method with maximum distance as the metric. The heatmap was generated using the pheatmap R package with the viridis color palette.

#### 4.2.2. Beta Regression Modeling for Tissue-Specific Isoform Detection

To identify tissue-specific relative isoform expression (Figure 2B), we implemented a beta regression analysis using the betareg R package (Cribari-Neto and Zeileis, 2010). The analysis was restricted to protein-coding transcripts from genes with at least two isoforms and a maximum expression of ≥ 1 TPM. For each transcript, relative expression was calculated as a proportion of the total expression of its parent gene. Beta regression models were fitted for each transcript with tissue as the primary predictor, while controlling for experimental batch as a covariate (relative_expression ∼ tissue + batch). P-values were adjusted for multiple testing using the Benjamini-Hochberg procedure, and transcripts with an FDR < 0.1 were considered significantly tissue-specific. For visualization, each significant transcript was assigned to the tissue in which it had the maximum beta coefficient.

#### 4.2.3. Functional Enrichment Analysis

Pathway enrichment analysis (Figure 2C) was conducted using the EnrichR (Xie et al., 2021) web service against the MSigDB Oncogenic Signatures (C5) gene set collection. For each tissue, a list of genes corresponding to the tissue-specific transcripts was submitted for analysis, using the complete set of tested transcripts as the background universe. A curated list of 27 relevant oncogenic pathways was selected for visualization. The combined enrichment score was calculated as the product of the -log10(adjusted p-value) and the z-score to capture both statistical significance and effect size.

#### 4.2.4. Isoform Abundance Differential Expression Analysis

To identify transcripts with tissue-specific or cell-line-specific absolute abundance, we performed differential expression analysis on the TPM values quantified by MPAQT (Apostolides et al., 2024). The analysis was conducted using the limma package in R (Ritchie et al., 2015), which is well-suited for complex experimental designs. A linear model was fitted for each transcript, with tissue or cell line as the primary predictor variable, while controlling for experimental batch effects (∼0 + tissue + batch or ∼ 0 + cell_line + batch). Differential expression was assessed using empirical Bayes moderated t-statistics. Transcripts were considered significantly tissue- or cell-line-specific based on an FDR-adjusted p-value threshold. Subsequent functional enrichment analyses on these gene lists were performed as described above.

#### 4.2.5. Ternary Simplex Analysis of Transcript Diversity

The ternary analysis in Figure 2D was performed following the ENCODE4 (Reese et al., 2023) methodology to decompose transcript diversity into contributions from alternative TSS, alternative TES, and alternative splicing. The analysis utilized the “inferred reference” transcript set from our Cerberus (Reese, 2023) annotation pipeline. For each gene, we quantified the number of unique TSSs (N_TSS), TESs (N_TES), and exon chains (N_EC). Splicing complexity was quantified using the ENCODE4 splicing ratio: (2 * N_EC) / (N_TSS + N_TES). These three components were normalized to sum to 1.0 to derive the ternary coordinates for each gene. The ternary plot was generated using the ggtern R package, with density visualized using hexagonal binning and a log10-transformed viridis color scale.

### 4.3. Training Data

We curated a comprehensive dataset of transcript isoforms by aggregating long-read RNA-seq data from multiple sources. This dataset includes 52 samples (two biological replicates for 26 cell lines) from the C2L2 project, along with 149 additional human and mouse samples from other long-read RNA-seq projects. These projects span diverse biological conditions, including normal tissues, various cancer types, COVID-19 studies, control groups, and drug-treated conditions. In total, the dataset comprises 201 human samples with 191,273 multi-exon transcripts, and 46 mouse samples with 106,453 multi-exon transcripts, each with a maximum standardized length of 65,536 nucleotides. The final dataset for Mach-1 training included 297,726 unique transcripts, providing over 7 billion single-nucleotide tokens annotated with species-specific identifiers (Table 1).

**Table 1.** Transcripts and tokens used to develop Mach-1.

| Species | Num. Transcripts | Num. Tokens |
| --- | --- | --- |
| human | 191,273 | 4,985,385,906 |
| mouse | 106,453 | 2,299,348,935 |
| total | 297,726 | 7,284,734,841 |

To assess Mach-1’s generalization performance, the data was split into training, validation, and test sets based on chromosome assignment (Table 2), with chromosome 10 reserved for validation and chromosome 8 designated as the test set.

**Table 2.** Training, validation, and test sets for Mach-1.

| Species | Split | Chromosome | Num. Transcripts | Num. Tokens | % of Tokens |
| --- | --- | --- | --- | --- | --- |
| human | train | 1-7,9,11-22,X,Y,M | 179,089 | 4,653,411,260 | 93.34 |
| human | validation | 10 | 6,169 | 177,889,685 | 3.57 |
| human | test | 8 | 6,015 | 154,084,961 | 3.09 |
| mouse | train | 1-7,9,11-19,X,Y | 96,504 | 2,084,601,665 | 90.66 |
| mouse | validation | 10 | 4,510 | 93,748,826 | 4.08 |
| mouse | test | 8 | 5,439 | 120,998,444 | 5.26 |

### 4.4. Model Architecture

#### 4.4.1. StripedHyena architecture

Mach-1 is based on StripedHyena (Poli et al., 2023b), an architecture for long-sequence language modeling (naW, b) that combines Hyena (Poli et al., 2023a) and rotary self-attention operators (Vaswani et al., 2017). The StripedHyena architecture can be described as a sequence of transformations applied to an input sequence *x* ∈ ℝ*^n^*^×^*^d^*, where *n* is the sequence length and *d* is the dimensionality of the model’s hidden states. Each layer (block) in StripedHyena is either a Hyena layer or a rotary self-attention layer, interleaved to balance the strengths of convolutional and attention mechanisms. Mach-1 consists of 16 blocks with a model width of 256 dimensions, where each block features sequence mixing and channel mixing layers to process information along the sequence and model width dimensions, respectively. Within the sequence mixing layers, Mach-1 utilizes 13 Hyena layers, alternated with 3 rotary self-attention layers at regular intervals. The channel mixing layers in Mach-1 use gated linear units. Additionally, Mach-1 applies root mean square layer normalization to the inputs of each layer.

#### 4.4.2. Hyena layers

The Hyena layers in Mach-1 are pivotal for efficiently capturing long-range dependencies in sequence data, offering a computationally efficient alternative to traditional attention mechanisms. The layers employ a blend of implicitly parametrized long convolutions and data-controlled gating functions, allowing them to process sequences with a computational complexity that scales sub-quadratically with the sequence length. While the self-attention mechanism captures global dependencies by computing pairwise interactions between all elements in a sequence—a powerful but computationally expensive approach with a complexity of *O*(*n*^2^*d*)—the Hyena mechanism achieves similar outcomes using a combination of implicit long convolutions and data-controlled gating mechanisms. These convolutions can be efficiently computed using the fast Fourier transform (FFT), allowing the Hyena mechanism to model dependencies over varying scales without explicit pairwise comparisons. This approach results in a more scalable complexity of *O(nd* log *n*), where *n* is the sequence length and *d* is the model width. Additionally, while self-attention requires explicit positional encodings to maintain the order of tokens in a sequence, Hyena uses a combination of positional encoding and windowed convolutions to encode positional information implicitly, offering greater flexibility in handling sequences of varying lengths. Detailed information of the architecture can be found in Poli et al. (Poli et al., 2023a).

#### 4.4.3. Self-Attention layers

Self-attention mechanisms are integral to Mach-1’s ability to model intricate dependencies between tokens in a sequence, regardless of their positions. Originally introduced in the Transformer architecture (Vaswani et al., 2017), self-attention computes the importance of each token in the sequence relative to others, enabling the model to focus on relevant parts of the sequence for a given task.

Given an input sequence *X*, self-attention operates by computing three matrices: queries *Q*, keys *K*, and values *V*:

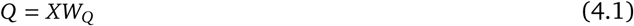

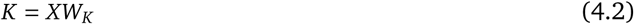

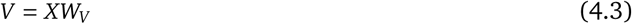

where *W_Q_*, *W_k_*, *W_V_* are learnable weight matrices, and *d_k_* is the dimensionality of the queries and keys. The attention scores are computed as:

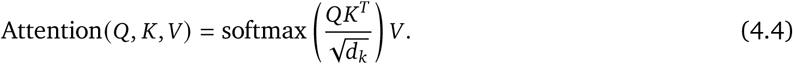

The softmax function ensures that the attention weights sum to one, allowing the model to distribute its focus across the sequence. The division by 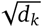 is a scaling factor that stabilizes gradients during training.

In Mach-1, the attention mechanism is enhanced by rotary position embeddings (RoPE) (Su et al., 2021), which encode positional information by rotating the query and key vectors:

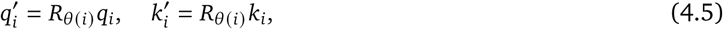

where *R_θ_*_(*i*)_ is a rotation angle that depends on the relative position of token *i* in the sequence. RoPE effectively captures relative positional relationships, allowing the model to distinguish between different token orders while maintaining position invariance.

### 4.5. Tokenization

In Mach-1, RNA sequences are tokenized at a single-nucleotide resolution, with an effective vocabulary of 16 tokens from a total of 32 characters. The tokenization process can be described mathematically as follows: Given an RNA sequence *s* = (*s*_1_*, …, s_n_*) of length *n*, each nucleotide *s_i_* is mapped to a token *t_i_* from the vocabulary *V*. The tokenization function *f* maps the sequence *s* to a sequence of tokens *t* = (*t*_1_*, …, t_n_*), where *t_i_* = *f* (*s_i_*).

For exonic sequences, the nucleotides are encoded using uppercase tokens (*A*, *C*, *G*, *U*), while intronic sequences use lowercase tokens (*a*, *c*, *g*, *u*). Additionally, special tokens (*W*, *X*, *Y*, *Z*) are used to encode non-RNA (DNA) sequences flanking the transcripts, representing nucleotides A, C, G, and T, respectively. Tokens *<*S*>* and *<*P*>* mark the transcription start site (TSS) and polyA site (PAS), respectively. The tokens *H* and *M* are used to specify whether the sequence belongs to a human or mouse, respectively.

### 4.6. Training Procedure

Mach-1 was pre-trained using causal language modeling (CLM) with a next token prediction (NTP) objective. This task involves predicting the next token in a sequence based on its preceding context, a process critical for learning the sequential dependencies inherent in RNA sequences. The training objective is to minimize the cross-entropy loss:

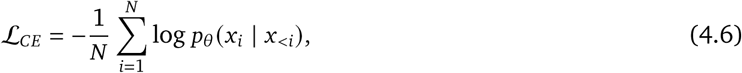

where *N* is the number of tokens in the sequence, *x_i_* is the true token at position *i*, and (*p_θ_ x_i_|x_<i_*)is the predicted probability of token *x_i_* given the input context.

From a probabilistic standpoint, minimizing the cross-entropy loss is equivalent to maximizing the likelihood of the observed data under the model, a principle known as Maximum Likelihood Estimation (MLE):

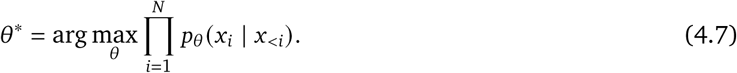

Taking the negative log of the likelihood gives us the cross-entropy loss. Perplexity, derived from cross-entropy, is a commonly used metric for evaluating language models:

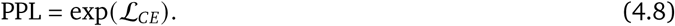

Perplexity can be interpreted as the exponential of the average log-loss per token, measuring the model’s uncertainty in predicting the next token.

### 4.7. Scaling Laws and Comparison with All-Hyena and All-Attention Architectures

In our study, we conducted a comprehensive scaling law analysis to compare the performance of various model architectures under a compute-optimized protocol. The primary objective was to determine the optimal balance between model size and dataset size to maximize the efficiency of computational resources. Our experiments explored model sizes ranging from 0.5 million to 40 million parameters, maintaining a fixed context length of 65,536 tokens. To ensure the reliability of our results, we gradually reduced the learning rate and repeated the experiments until convergence was achieved. Model performance was evaluated using the perplexity metric.

To carry out the scaling law analysis, we defined a series of compute budgets ranging from 10^15^ to 10^19^ FLOPs. For each budget, we calculated the FLOPs required to process a fixed input size, representing the cost of running the model. We then identified the compute-optimal allocation by selecting a range of model sizes and determining the number of training tokens needed to fully utilize the compute budget. To pinpoint the optimal compute allocation, we fitted a second-order polynomial to the relationship between log model size and perplexity. The minimum point of this function indicates the compute-optimal point, which signifies the most efficient allocation of model size and training tokens for a given compute budget.

To further evaluate performance, after identifying the optimal architecture—comprising 13 Hyena layers interleaved with 3 attention layers (16 layers total)—we compared it against two other configurations: an All-Hyena model (16 Hyena layers, similar to HyenaDNA) and an All-Attention model (16 attention layers, similar to GPT). All models were trained using the same hyperparameters and were allowed to train until the validation loss reached a plateau.

#### 4.7.1. Motif Discovery Across PAS Embeddings Clusters

Mach-1 is composed of 16 layers, with each layer progressively encapsulating more complex and abstract information from the input sequences. As the model deepens, it learns increasingly sophisticated concepts, patterns, and dependencies. To understand the type of information encoded within each layer, the embeddings from different layers can be analyzed to observe how the samples (transcripts) are distributed across these embeddings in the hidden space. Specifically, we focused on the penultimate layer’s embeddings of the PAS token for each input sequence, using these embeddings as representations of the entire sequence.

To identify distinct patterns within these embeddings, we performed k-means clustering on the first 50 principal components of the embeddings, using a range of cluster numbers to identify the optimal number of clusters (by maximizing the silhouette value)—we found that the PAS embeddings were best described by 18 distinct clusters. To examine whether these clusters correspond to distinct sequence features, we used *MEME-ChIP* (Machanick and Bailey, 2011) in the differential enrichment mode to identify motifs that were significantly enriched in each cluster compared to sequences of all other clusters, along with their localization profiles relative to the poly(A) site.

#### 4.7.2. Splice Sites and Rbfox1 Motifs Likelihood Analysis

Mach-1 was trained using a next-token (nucleotide) prediction task, leveraging causal language modeling to develop a deep understanding of the biological language of pre-mRNA transcripts. This approach enables Mach-1 to become a splice-aware RNA foundation model, capable of generating sequence embeddings and estimating their likelihoods. To evaluate how effectively the model understands regulatory elements, an *in silico* mutational scanning experiment was conducted to identify which input tokens (nucleotides) are critical for the model’s likelihood estimation.

In this experiment, each token in the sequence was mutated into all possible alternatives, resulting in seven mutated sequences plus the original reference sequence—eight sequences in total. Mach-1 then estimated the likelihood of each sequence. These likelihoods were normalized into a categorical probability distribution using the softmax function, where each element of the distribution represents the probability of the corresponding sequence. Since the only difference between the sequences is one token, the resulting probability distribution indicates the importance of the reference nucleotide to the model. A higher probability for the sequence with the reference token suggests that this nucleotide is critical for the model’s predictions.

This experiment was performed on all tokens across all transcripts of the Grin1 gene in the mouse genome. Grin1 was chosen because previous research (Gehman et al., 2011) had investigated Rbfox1 motifs located downstream of one of the cassette exons in Grin1 transcript isoforms. The analysis revealed that Mach-1 assigns high likelihood scores to splice site signals across the gene. Furthermore, when examining the +8 and +262 tokens downstream of the exon reported in the referenced study, it was observed that the model recognizes the regulatory significance of Rbfox1 motifs, both in proximity to and at a distance from the cassette exon of interest. This demonstrates the model’s capability to understand and predict key regulatory elements within transcript sequences.

#### 4.7.3. Systematic Analysis of Mach-1 Sensitivity to Splice Site Mutations

To systematically assess Mach-1’s sensitivity to splice site mutations, we first identified cassette exon events using *SUPPA2* (Trincado et al., 2018) and selected representative isoforms that support the inclusion or exclusion of each event. For each cassette exon, we chose the isoforms with the highest coverage—one representing inclusion and one representing exclusion. For each representative isoform, we then removed (replaced with ‘N’) the splice site signal (two nucleotides in the intronic region) upstream or downstream of the cassette exon, representing acceptor or donor sites, respectively, and calculated the corresponding Δlog-Likelihood. For all donor or acceptor splice sites, we obtained paired Δlog-Likelihoods for isoforms supporting cassette exon inclusion and those supporting exclusion. These pairs were plotted as points in scatter plots of Δlog-Likelihood in transcripts excluding versus including the exon (Figure 4B). The statistical significance of the differences in Δlog-Likelihoods between exon-inclusion and exon-exclusion isoforms was assessed using the one-sided Wilcoxon signed-rank test.

#### 4.7.4. Systematic Analysis of Mach-1 Sensitivity to RBP Binding Sites

We obtained peak calling BED files (narrowPeak format) for 150 RNA-binding proteins (RBPs) in two cell lines, K562 and HepG2. The list of RBPs includes: AARS, AATF, ABCF1, AGGF1, AKAP1, AKAP8L, APOBEC3C, AQR, BCCIP, BCLAF1, BUD13, CDC40, CPEB4, CPSF6, CSTF2, CSTF2T, DDX21, DDX24, DDX3X, DDX42, DDX51, DDX52, DDX55, DDX59, DDX6, DGCR8, DHX30, DKC1, DROSHA, EFTUD2, EIF3D, EIF3G, EIF3H, EIF4G2, EWSR1, EXOSC5, FAM120A, FASTKD2, FKBP4, FMR1, FTO, FUBP3, FUS, FXR1, FXR2, G3BP1, GEMIN5, GNL3, GPKOW, GRSF1, GRWD1, GTF2F1, HLTF, HNRNPA1, HNRNPC, HNRNPK, HNRNPL, HNRNPM, HNRNPU, HNRNPUL1, IGF2BP1, IGF2BP2, IGF2BP3, ILF3, KHDRBS1, KHSRP, LARP4, LARP7, LIN28B, LSM11, MATR3, METAP2, MTPAP, NCBP2, NIP7, NIPBL, NKRF, NOL12, NOLC1, NONO, NPM1, NSUN2, PABPC4, PABPN1, PCBP1, PCBP2, PHF6, POLR2G, PPIG, PPIL4, PRPF4, PRPF8, PTBP1, PUM1, PUM2, PUS1, QKI, RBFOX2, RBM15, RBM22, RBM5, RPS11, RPS3, SAFB, SAFB2, SBDS, SDAD1, SERBP1, SF3A3, SF3B1, SF3B4, SFPQ, SLBP, SLTM, SMNDC1, SND1, SRSF1, SRSF7, SRSF9, SSB, STAU2, SUB1, SUGP2, SUPV3L1, TAF15, TARDBP, TBRG4, TIA1, TIAL1, TRA2A, TROVE2, U2AF1, U2AF2, UCHL5, UPF1, UTP18, UTP3, WDR3, WDR43, WRN, XPO5, XRCC6, XRN2, YBX3, YWHAG, ZC3H11A, ZC3H8, ZNF622, ZNF800, ZRANB2.

Using *SUPPA2*, we identified cassette exon events in our inferred transcriptome and selected the isoform with the highest expression in either K562 or HepG2 (depending on the cell line used in the narrowPeak data) to represent exon inclusion for each cassette exon. We then determined the overlap between RBP-bound regions and the regions of cassette exons, including their flanking introns but excluding splice sites and ±5 flanking nucleotides. To quantify the impact of RBPs on cassette exon inclusion, we mutated (replaced with ‘N’) each RBP-bound region, one at a time, within the representative isoform and calculated the Δlog-Likelihood between the mutant and reference (unaltered) isoform sequences. To serve as a negative control, we also randomly selected non-RBP regions of similar length distribution that did not overlap with any RBP-bound regions. These randomly chosen regions were mutated in a similar fashion, and the Δlog-Likelihood was calculated. For each RBP and non-RBP control region, we generated boxplots of Δlog-Likelihoods. We then sorted RBPs based on their median Δlog-Likelihood, showing that RBPs with a higher median Δlog-Likelihood were more significantly enriched in the splicing gene set (splicing factors), indicating their crucial role in splicing regulation.

#### 4.7.5. Experimental Validation of RBP-Mediated Splicing Predictions

To experimentally validate Mach-1’s predictions regarding the functional impact of RBP binding sites, we selected three RBPs (TAF15, SAFB, and SAFB2) for knockdown experiments in K562 cells using CRISPR interference (CRISPRi). Single-guide RNA (sgRNA) sequences for CRISPRi-mediated gene knockdown were cloned into pCRISPRia-v2 (Addgene #84832) via BstXI-BlpI sites. The cloned plasmid was co-transfected with plasmids pMD.2G and pCMV-dR8.91 using TransIT-Lenti (Mirus) into HEK293T cells according to the manufacturer’s manual. Viral supernatant was collected 48 hours after transfection and added to K562 cells expressing a Zim3-KRAB-dCas9 fusion protein (Addgene #154472) in the presence of 8 *μ*g mL^−1^ polybrene (Millipore). After 48 hours post-transduction, cells were selected in the presence of 8 *μ*g mL^−1^ puromycin. The single-guide RNA sequences were as follows:

sgControl: GACCAGGATGGGCACCACCC

sgTAF15: GGGCGGCCGGAGCTGTACTG

sgSAFB: GCGCGCTGGGGCGACTGGAG

sgSAFB2: GCGGCGGTGTGCGACTGAGT

Following selection, RNA was extracted from the knockdown and control cell populations and subjected to both short- and long-read RNA sequencing. To quantify splicing changes, we used SUPPA2 to calculate the change in Percent Spliced-In (ΔPSI) for cassette exon events between each knockdown and the control condition. P-values were adjusted using the Benjamini-Hochberg method. Our analysis focused on cassette exons where the flanking intronic regions (±256 bp, excluding ±5 bp from the splice site) contained binding motifs for the RBP that was knocked down. The experimentally derived ΔPSI values for significant events (FDR < 0.05) were then compared to the Δlog-likelihood (ΔLL) scores previously calculated by Mach-1 for the in silico mutation of the corresponding RBP binding sites. The relationship between ΔPSI and ΔLL was assessed using both Pearson and Spearman correlation analyses. Additionally, we evaluated the correlation between the ΔLL scores and the log-fold change (logFC) in the expression of the transcripts hosting these cassette exons.

#### 4.7.6. Discovery of Sequence Features (Motifs) to which Mach-1 is Sensitive

For all identified cassette exon events, we focused on the cassette exon itself along with 256 bp of its flanking regions (both upstream and downstream), while excluding the splice site signals and the ±5 nucleotides surrounding them. We chose 256 bp as the flanking region length because it was computationally feasible while providing sufficient context for analysis. To determine which features contribute most significantly to cassette exon inclusion, we selected the isoform with the highest expression supporting exon inclusion as the representative isoform. We used a sliding window approach to systematically analyze the specific regions of interest. A window size of 20 nucleotides with a stride of 10 nucleotides was applied across the target regions. For each window, the content was replaced with ‘N’, and we calculated the Δlog-Likelihood between the mutant isoform (altered) and the reference isoform (unaltered). In total, we analyzed over 250,000 windows with corresponding Δlog-Likelihood values. We sorted the regions based on Δlog-Likelihood and selected the top 50% (those with Δlog-Likelihood values above the median) for further analysis. We then used the MEME suite to perform motif discovery within these high-impact regions. This analysis identified eight enriched sequence motifs, three of which were found to match known RBP binding sites using the Tomtom tool in conjunction with the Ray2013 dataset (Ray et al., 2013).

### 4.8. Zero-shot Prediction Tasks

#### 4.8.1. Isoform Absolute Abundance

To assess Mach-1’s zero-shot prediction performance for isoform absolute abundance, we provided the entire pre-mRNA sequence of each transcript isoform into Mach-1 and calculated two likelihood-based metrics: (1) the log-likelihood of the entire pre-mRNA transcript sequence, and (2) the log-likelihood of the last token (polyadenylation site, denoted as “P”). Both metrics were evaluated, and the one yielding the stronger correlation with measured isoform abundances was reported.

For causal language models (CLMs) such as Mach-1, Evo 2 (Brixi et al., 2025), and HyenaDNA (Nguyen et al., 2023), likelihoods are computed directly by conditioning on preceding nucleotides. Mach-1 processes full pre-mRNA sequences (64K nucleotide context), while Evo 2 and HyenaDNA operate on mature mRNA sequences with 1M context windows. Each model produced both sequence-wide and final-token likelihoods, and we reported the best-performing metric. A summary of abundance-prediction evaluation procedures across all models is provided in Table 3.

**Table 3.** Zero-shot Isoform Abundance Prediction Tasks.

| Model | Context (nt) | Type | How |
| --- | --- | --- | --- |
| Mach-1 | 64K | CLM | $LL(\text{EntirePre-mRNATranscriptSequence})$;<br>$LL(\text{LastTokenEntirePre-mRNATranscriptSequence})$ |
| Evo-2 | 1M | CLM | $LL(\text{EntireMature-mRNATranscriptSequence})$;<br>$LL(\text{LastTokenEntireMature-mRNATranscriptSequence})$ |
| HyenaDNA | 1M | CLM | $LL(\text{EntireMature-mRNATranscriptSequence})$;<br>$LL(\text{LastTokenEntireMature-mRNATranscriptSequence})$ |
| Nucleotide Transformer v2 | ~12K | MLM | If fits: $PLL(\text{EntireMature-mRNATranscriptSequence})$; if not:<br>$AvgPLL(\text{ChunkedMature-mRNATranscriptSequence})$ |
| DNABERT-2/S | ~2K | MLM | If fits: $PLL(\text{EntireMature-mRNATranscriptSequence})$; if not:<br>$AvgPLL(\text{ChunkedMature-mRNATranscriptSequence})$ |
| RNA-FM | 1024 | MLM | If fits: $PLL(\text{EntireMature-mRNATranscriptSequence})$; if not:<br>$AvgPLL(\text{ChunkedMature-mRNATranscriptSequence})$ |
| Borzoi | 524K | CNN+Attention | $\text{LogCov} = \log(\text{Avg}(\text{EntirePre-mRNAExonsCov}))$;<br>$\text{LogCov} = \log(\text{Avg}(\text{UniqueRegionsOfPre-mRNAExonsCov}))$; averaged over 4 models |
| Pangolin | 10K | Score | $\text{Avg}(\text{EntirePre-mRNA SpliceJunctionVariantScores})$; up to 8 variants per intron (2 exonic + 2 intronic at both 5 and 3 splice sites) |
| GPN-MSA | 128 | Score | $\text{Avg}(\text{EntirePre-mRNA SpliceJunctionVariantScores})$ |
| CADD v1.7 | 500 | Score | $\text{Avg}(\text{EntirePre-mRNA SpliceJunctionVariantScores})$ |
| SpliceAI | 10K | Score | $\text{Avg}(\text{EntirePre-mRNA SpliceJunctionVariantScores})$ |

Masked language models (MLMs), including Nucleotide Transformer v2 (Dalla-Torre et al., 2025), DNABERT2 (Zhou et al., 2023), and RNA-FM (Shen et al., 2024), were evaluated using the pseudo-log-likelihood (PLL) method (Salazar et al., 2020). Because MLMs predict masked tokens based on surrounding context, PLL for a sequence was computed by iteratively masking each nucleotide and summing the log-probabilities of predicting it correctly. When the entire mature mRNA sequence fit within the model’s context window (e.g., 12K for Nucleotide Transformer v2, 2K for DNABERT-2/S, and 1K for RNA-FM), we used the full-sequence PLL; otherwise, we computed the average PLL across overlapping sequence chunks.

For the Borzoi model (Linder et al., 2025), which combines convolutional and attention-based layers, we estimated abundance using coverage-derived metrics. Two variants were evaluated: (1) the logarithm of the average exon coverage across the entire pre-mRNA transcript sequence, and (2) the logarithm of the average coverage of its unique exonic regions. Each was averaged across four trained Borzoi models, and we reported the metric achieving the stronger correlation with experimental abundance.

Score-based models such as Pangolin (Zeng and Li, 2022), GPN-MSA (Benegas et al., 2024), CADD v1.7 (Rentzsch et al., 2019), and SpliceAI (Jaganathan et al., 2019) were evaluated by averaging their variant scores across all splice junctions in each pre-mRNA. For each tool, this includes up to eight variants per intron (two exonic and two intronic positions for both 5^′^ and 3^′^ splice sites).

#### 4.8.2. Trapped Exons Abundance Task

In exon-trapping assays, a fixed vector backbone is used to test whether inserted genomic fragments can function as exons. We used the same vector sequence as Stepankiw et al. (2023), who screened approximately 1.25 million genomic fragments for exon-trapping potential and reported their measured inclusion abundances. The evaluation procedure for all models in this task is summarized in Table 4.

**Table 4.** Zero-shot Exon Trap Prediction Tasks. Models evaluated on vector-based exon-trapping assays following Stepankiw et al. (2023). Both exon and intron configurations were scored, and the better-performing metric was reported.

| Model | Context (nt) | Type | How |
| --- | --- | --- | --- |
| Mach-1 | 64K | CLM | $\Delta LL = LL(\text{Vector} + \text{TrappedExon}) - LL(\text{Vector} + \text{Intron})$ |
| Evo-2 | 1M | CLM | $\Delta \text{AvgLL} = \text{AvgLL}(\text{VectorWithTrappedExon}) - \text{AvgLL}(\text{VectorWithoutTrappedExon})$ |
| HyenaDNA | 1M | CLM | $\Delta \text{AvgLL} = \text{AvgLL}(\text{VectorWithTrappedExon}) - \text{AvgLL}(\text{VectorWithoutTrappedExon})$ |
| Nucleotide Transformer v2 | ~12K | MLM | $\Delta \text{AvgPLL} = \text{AvgPLL}(\text{VectorWithTrappedExon}) - \text{AvgPLL}(\text{VectorWithoutTrappedExon})$ |
| DNABERT-2/S | ~2K | MLM | $\Delta \text{AvgPLL} = \text{AvgPLL}(\text{VectorWithTrappedExon}) - \text{AvgPLL}(\text{VectorWithoutTrappedExon})$ |
| RNA-FM | 1024 | MLM | $\Delta \text{AvgPLL} = \text{AvgPLL}(\text{VectorWithTrappedExon}) - \text{AvgPLL}(\text{VectorWithoutTrappedExon})$ |
| Borzoi | 524K | CNN+Attention | $\text{PSI} = \text{Avg}(\text{Cov}(\text{TrappedExons})) / \text{Avg}(\text{Cov}(\text{AllExons})); \Delta \log \text{PSI} = \log(\text{PSI} / (1 - \text{PSI}));$<br>averaged over 4 models |

For each fragment, two sequences were generated: one with the fragment inserted as an intron (lowercase, e.g., acgt) and one as an exon (uppercase, e.g., ACGT). Both full vector sequences were input into Mach-1, and we calculated two likelihood-based metrics: (1) the log-likelihood difference of the entire vector sequence with and without the exon, and (2) the difference in last-token likelihoods between the two sequences. The better-performing metric was reported. Formally, the main sequence-wide difference is:

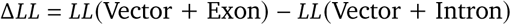

which serves as a proxy for the likelihood of exon inclusion. The resulting Δ*LL* values were compared against measured exon-trapping abundances.

For Evo 2 and HyenaDNA, we computed both total and average log-likelihood differences (AvgLL) between the exon (the trapped exon sequence is included) and intron (the trapped exon sequence is excluded) configurations and reported the best-performing metric. For MLMs such as Nucleotide Transformer v2, DNABERT-2, and RNA-FM, we computed the difference in average pseudo-log-likelihood (AvgPLL) between the two sequences and used either the full-sequence or chunked form depending on fit; the better one was reported. For Borzoi, we computed the percent-spliced-in (PSI) as the ratio between the average coverage of trapped exons and all exons in the vector, and derived ΔlogPSI = log(PSI/1 – PSI). Two PSI-based metrics, analogous to those in the abundance task, were calculated and averaged over four trained Borzoi models, with the stronger-performing metric reported.

Variant-score models such as Pangolin, GPN-MSA, CADD, SpliceAI, and AlphaGenome (Avsec et al., 2025) were not included in this task because they produce variant-level rather than sequence-level predictions, and thus cannot distinguish exon versus intron configurations.

#### 4.8.3. Pathogenicity Prediction for Non-coding Variants

To evaluate Mach-1’s ability to predict the impact of non-coding mutations, we performed a zero-shot pathogenicity classification using variants from ClinVar (Landrum et al., 2014) and SpliceVarDB. Variants were labeled as pathogenic or benign according to their curated clinical annotations. For evaluation, “pathogenic” and “likely pathogenic” variants were merged into a single class, while “benign”, “likely benign”, and “VUS” were merged into another, forming a binary classification (pathogenic vs. non-pathogenic). The variant-effect prediction setup is summarized in Table 5.

**Table 5.** Zero-shot Variant Effect Prediction Tasks.

| Model | Context (nt) | Type | How |
| --- | --- | --- | --- |
| Mach-1 | 64K | CLM | $\Delta LL = LL(\text{EntireMutantTranscriptSequence}) - LL(\text{EntireWildTranscriptSequence})$ |
| Evo-2 | 1M | CLM | $\Delta LL = LL(8\text{kMutantGenomeSequence}) - LL(8\text{kRefGenomeSequence})$ |
| HyenaDNA | 1M | CLM | $\Delta LL = LL(8\text{kMutantGenomeSequence}) - LL(8\text{kRefGenomeSequence})$ |
| Nucleotide Transformer v2 | ~12K | MLM | $\text{Pseudo}\Delta LL = \text{PLL}(8\text{kMutantGenomeSequence}) - \text{PLL}(8\text{kRefGenomeSequence})$ |
| DNABERT-2/S | ~2K | MLM | $\text{Pseudo}\Delta LL = \text{PLL}(8\text{kMutantGenomeSequence}) - \text{PLL}(8\text{kRefGenomeSequence})$ |
| RNA-FM | 1024 | MLM | $\text{Pseudo}\Delta LL = \text{PLL}(8\text{kMutantGenomeSequence}) - \text{PLL}(8\text{kRefGenomeSequence})$ |
| Pangolin | 10K | Score | Variant Score |
| GPN-MSA | 128 | Score | Variant Score |
| CADD v1.7 | 500 | Score | Variant Score |
| SpliceAI | 10K | Score | Variant Score |

For each variant, two transcript sequences—reference and mutant—were constructed. We computed two Mach-1-based metrics: (1) the difference in total sequence log-likelihood, and (2) the difference in last-token likelihood. The better-performing of these two Δlog-likelihoods was used for downstream classification:

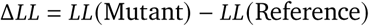

A more negative Δ*LL* indicates a mutation that decreases transcript likelihood and is thus more likely to be deleterious.

For Evo 2 and HyenaDNA, Δ*LL* was calculated over 8k genome-centered sequences, and both total and average log-likelihood differences were tested, reporting the best. For MLMs (Nucleotide Transformer v2, DNABERT-2, RNA-FM), we computed the difference in pseudo-log-likelihood (PseudoΔLL) between 8k genome-centered mutant and reference sequences.

The Borzoi model was excluded from ClinVar and SpliceVarDB variant-effect analyses, as its original study (Linder et al., 2025) did not benchmark variant pathogenicity or provide mutation-level prediction outputs.

For score-based models such as Pangolin, GPN-MSA, CADD v1.7, and SpliceAI, we used their provided variant scores for classification, while variant scores from AlphaGenome were obtained through its public API. Models with highly similar architectures and training datasets—such as ERNIE and RiNALmo—were excluded from all analyses to avoid redundancy with previously benchmarked models.

#### 4.8.4. Assessing Somatic Mutation Effects on Overall Isoform Expression in a Zero-Shot Scenario

To evaluate Mach-1’s capacity to predict the impact of somatic mutations on transcriptome architecture, we utilized cell line-specific mutation data from the DepMap database. Our analysis focused on assessing the model’s ability to predict mutation-induced changes in transcript expression and alternative splicing patterns without additional cell line-specific training, implementing a zero-shot prediction framework. Initially, we employed SUPPA2 to identify differentially spliced exons between cell line pairs within each tissue type, concentrating our analysis on splicing events near the somatic mutation sites. The identified novel alternative splicing events were categorized into five distinct types: alternative 3’ splice site (A3), alternative 5’ splice site (A5), alternative first exon (AF), alternative last exon (AL), and retained intron (RI). For each identified mutation, we computed the normalized change in isoform likelihood (ΔLL) between reference and mutant sequences using Mach-1. To validate these predictions, we quantified changes in exon inclusion levels (ΔPSI). The relationship between ΔLL and ΔPSI was evaluated using Pearson correlation analysis.

To further validate Mach-1’s ability to predict the functional consequences of somatic mutations, we extended our analysis to overall isoform expression changes. We utilized the DepMap database to systematically identify somatic mutations that were present in a subset of our C2L2 cell lines but absent in other cell lines from the same tissue lineage. For this set of lineage-specific mutations, we focused on the transcripts directly affected by each variant. The ground truth for the mutational impact was established by calculating the log-fold change (logFC) in expression for each affected transcript between the cell lines harboring the mutation and those that did not. In parallel, we used Mach-1 in a zero-shot framework to predict the impact of each mutation by computing the change in log-likelihood (ΔLL) between the reference (wild-type) and mutant transcript sequences. Finally, the relationship between the predicted ΔLL values and the observed logFC in expression was evaluated using Pearson correlation analysis to determine the model’s predictive accuracy.

#### 4.8.5. CRISPR mutation and expression (band) analysis

For SNP editing, crRNA, tracrRNA, and ssODN HDR templates were designed and synthesized by Integrated DNA Technologies (IDT). Cas9 ribonucleoprotein (RNP) complexes were assembled and nucleofected into HEK293 cells using the Lonza SF Cell Line 96-well Nucleofector® Kit according to the CM-130 protocol, with 0.5 × 10^6 cells per electroporation. Cells were harvested 72 h post-electroporation for RNA and genomic DNA extraction. For splicing analysis, cDNA was synthesized using the Maxima H Minus First Strand cDNA Synthesis Kit (Thermo Fisher Scientific), followed by gene-specific PCR and visualization on a 2% agarose gel. To confirm editing efficiency, PCR was performed with primers flanking the targeted loci, and the amplicons were subjected to Nanopore sequencing (Plasmidsaurus).

#### 4.8.6. Predicting Isoform Abundance Using Embeddings

To assess the utility of embeddings generated by Mach-1 and other genomic foundation models, we trained downstream cell line-specific models to predict isoform abundance. For each model (Mach-1, Borzoi, Evo2, DNABERT-2, Nucleotide Transformer, HyenaDNA, RNAFM, and SpliceAI), we generated a feature representation for each isoform by averaging the last hidden-layer embeddings across the entire transcript sequence. For models with sufficient context length (Mach-1, Borzoi, Evo2, and HyenaDNA), the full pre-mRNA sequence served as input. For models with shorter context windows (DNABERT-2, Nucleotide Transformer, RNAFM, and SpliceAI), the mature-mRNA sequence was used instead.

The abundance data was obtained using MPAQT (Apostolides et al., 2024), which integrated long-read and short-read RNA-seq data from the C2L2 dataset, with the log-transformed TPM (transcripts per million) values serving as the prediction target. For each cell line, a separate gradient boosting regressor was trained using the scikit-learn library, with the model-specific embeddings as the feature set and the observed log-TPM values as the target. The regressors were trained with a data split strategy involving all chromosomes except 10 and 8 for training, chromosome 10 for validation, and chromosome 8 for testing. Performance was evaluated by calculating the Pearson correlation coefficient between the predicted and observed isoform abundance values for each cell line on the test set (chromosome 8) isoforms.

#### 4.8.7. Predicting Differential Isoform Expression Using Embeddings

To evaluate the ability of various model embeddings to predict differential isoform expression, we conducted DE analysis between pairs of cell lines within each tissue type. For each pair (e.g., cell line A vs. cell line B), we identified differentially expressed transcripts using LIMMA (Ritchie et al., 2015) with long-read RNA-seq data, labeling isoforms as positive if they were significantly upregulated (adjusted *p*-value < 0.05 and log2FC > 1) in cell line A. We then formulated a binary classification task to predict this upregulation using embeddings from Mach-1 and all other benchmarked models.

For each model, the feature set consisted of the averaged last-layer embeddings for each transcript. As before, embeddings for Mach-1, Borzoi, Evo 2, and HyenaDNA were derived from full pre-mRNA sequences, while embeddings for DNABERT-2, Nucleotide Transformer, RNA-FM, and SpliceAI were derived from mature-mRNA sequences due to context length limitations. It is important to note that these classification tasks were not symmetric—predicting upregulation in cell line A versus B is a distinct task from the reverse. For each task, we trained a random forest classifier using scikit-learn, employing a data split that reserved chromosome 10 for validation and chromosome 8 for testing. The performance of each model was evaluated by calculating the Area Under the Receiver Operating Characteristic (AUROC) and the Area Under the Precision-Recall Curve (AUPRC) on the test set isoforms.

#### 4.8.8. Transcript Sequence Generation

To generate transcript sequences, we utilized a causal language modeling approach with the Mach-1 model. Causal language models operate in an autoregressive manner, predicting the next token in a sequence based on the preceding context. This autoregressive nature allows these models to generate sequences in a coherent, step-by-step manner, maintaining both logical and semantic consistency throughout the sequence.

In our workflow, sequences were sampled from Mach-1 using standard techniques for autoregressive models, such as top-k sampling and temperature-based decoding. These methods help strike a balance between exploration and exploitation, allowing the model to produce diverse yet meaningful outputs. Mach-1 particularly benefits from the fast recurrent mode of its Hyena layers, which enhances computational efficiency by reducing both latency and memory requirements during sequence generation (naW, a).

To guide the sequence generation process, we employed standard language model prompting techniques. This involved conditioning the model with a given prefix, which served as the initial context for generating the subsequent output. For species-conditional generation, we used a single token representing the desired species (H for human or M for mouse) as the prompt, enabling the model to generate transcript sequences specific to the chosen species.

To compare the generated sequences to the reference transcriptome, we performed several analyses. We compared the distributions of sequence lengths, exon/intron counts, and exon/intron lengths. K-mer frequency correlations (for k=1 to 8) were calculated for exonic and intronic regions separately using both Pearson and Spearman methods. Motif enrichment analysis was performed separately on the reference and generated sequences using the MEME suite in differential enrichment mode, with statistical significance determined by MEME’s E-values. Enriched motifs identified in the two sets were then compared using Tomtom, which measures similarity between motif position weight matrices and reports significance using q-values (FDR-adjusted p-values). SpliceAI was used to score the donor and acceptor splice sites in both sets.

#### 4.8.9. Codon Usage Analysis of Generated Sequences

To evaluate the codon usage of sequences generated by Mach-1, we prompted the model to generate over 50,000 human transcript-like sequences. Each generated sequence contained intronic regions represented in lowercase. To identify open reading frames (ORFs) within these sequences, we first performed in-silico splicing by removing the intronic (lowercase) regions. After splicing, we utilized the *orfipy* package in Python (Singh and Wurtele, 2021) to identify ORFs within the sequences. The lengths of these ORFs were compared between the generated and reference sets. We then extracted the codons from the identified ORFs and analyzed their usage within the generated sequence dataset. Codon usage was compared to that in human reference transcripts to assess how closely the generated sequences resembled natural human transcripts.

#### 4.8.10. RNA sequence synthesis and sequencing

Designed RNA sequences were synthesized by Twist Bioscience and cloned into the pTwist CMV Puro backbone. The pooled plasmid library was transfected into HEK293 cells using TransIT®-Lenti Transfection Reagent, and total RNA was harvested 48 h post-transfection using the Zymo Quick-RNA Microprep Kit. First-strand cDNA was synthesized with the Maxima H Minus First Strand cDNA Synthesis Kit (Thermo Fisher Scientific), and amplicon libraries were generated using Phusion High-Fidelity Master Mix (New England Biolabs). Illumina sequencing adapters were added in a second round of PCR amplification, and the resulting libraries were subjected to next-generation sequencing according to standard Illumina protocols

## 5. Data availability

Raw short- and long-read RNA-sequencing data generated in this study are available at GEO: GEO accession number GSE280041, SRA accession number SRP540164.

## 6. Code and model availability

We make code and tools for model exploration available at the following links:

- Mach-1 model repository: https://github.com/goodarzilab/mach-1
- Mach-1 manuscript repository (notebooks, processed data, figures): https://github.com/csglab/mach-1-manuscript

## 7. Acknowledgments

This work was supported by grants from the Canadian Institutes of Health Research (CIHR) [PJT-173317] to HSN, Natural Sciences and Engineering Research Council of Canada (NSERC) [RGPIN-2019-04460] and the Canada Foundation for Innovation (CFI) JELF [project 40781] to AE, support by the Arc Institute to HG, and resources provided by Calcul Québec (calculquebec.ca) and the Digital Research Alliance of Canada (alliance-can.ca) to HSN and AE. HSN holds a CIHR Canada Research Chair. HG is an Arc Institute Core Investigator. Research in the lab of HG is supported by the Arc Institute. AS holds a Doctoral Training Scholarship from Fonds de recherche du Québec – Nature et technologies (FRQNT).

## 8. Author contributions

A.S., H.G., H.S.N., and A.E. conceived the project. H.G., H.S.N., A.E., and V.R. supervised the project. A.S. performed computational preprocessing of the C2L2 long- and short-read data, including pipeline development and QC. B.C. led experimental data generation for the C2L2 compendium, including cell line preparation and sequencing. S.W. assisted with sample preparation. S.Z. contributed to CRISPR mutation and synthetic transcript experiments. A.S. designed, implemented, and conducted all model training. M.N. contributed to model training and evaluation. A.S. performed model evaluation across all benchmarks. M.N. assisted with generation-based evaluations. N.B. assisted with fine-tuned abundance prediction evaluations. A.P. assisted with tissue- and cell line–specific isoform usage analyses. A.H.-C. assisted with differential expression analyses of the C2L2 cell lines. R.D. curated and prepared DepMap mutation data for isoform impact analysis. A.M.N. assisted with visualizations of model outputs and analyses. B.C. performed knockdown-based validation experiments. S.Z. contributed to experimental validations with CRISPR mutations and synthetic transcript generation. A.S. and B.C. wrote the first draft of the manuscript with input from all co-authors. All authors revised and approved the final manuscript.

## Supplementary Material

### A. Supplementary figures

**Figure S1.**
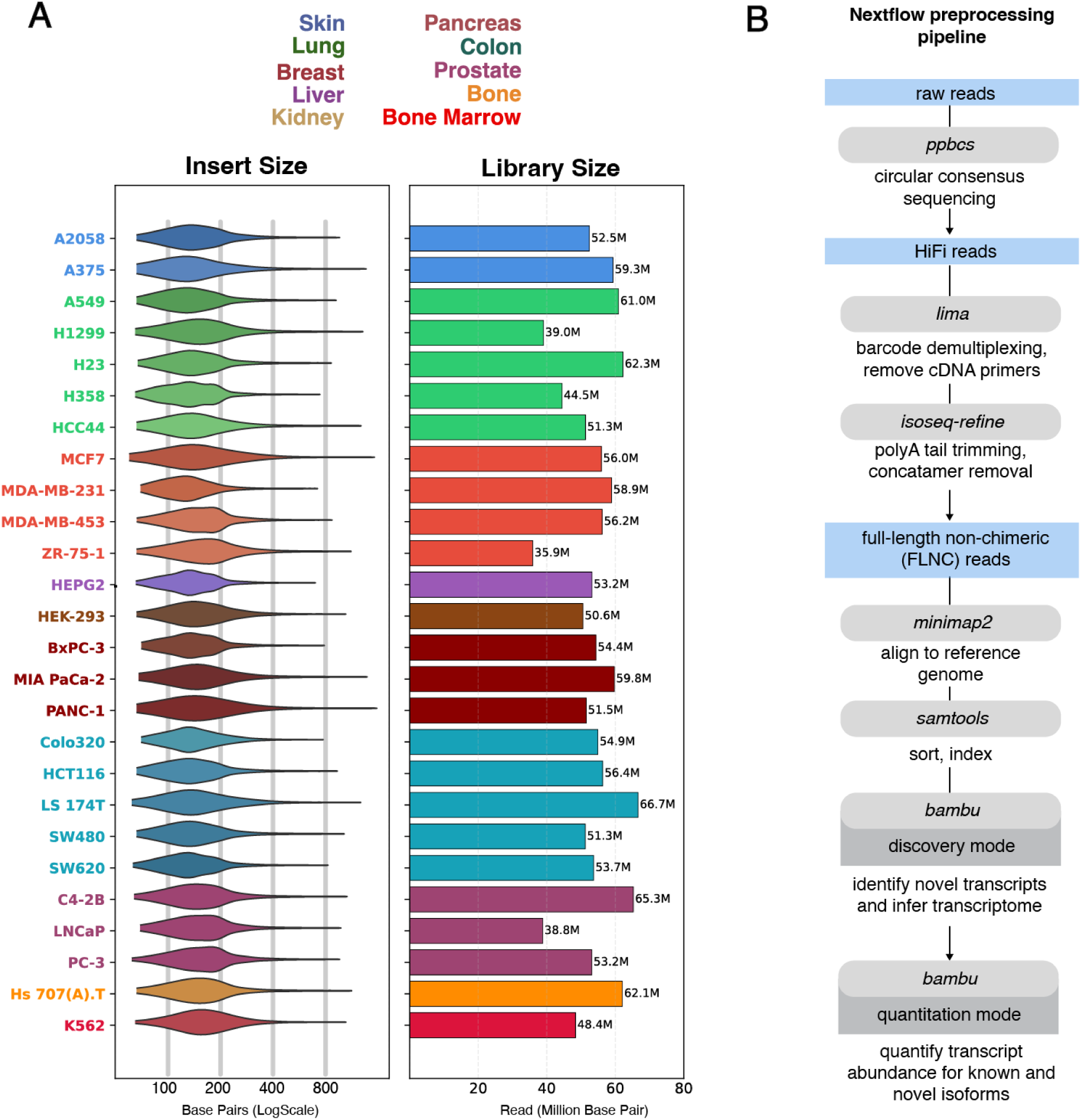
Long-read RNA-seq data processing pipeline and summary statistics for matched short-read RNA-seq data. (**A**) Summary statistics for the matched short-read RNA sequencing libraries for each of the 26 cell lines in the C2L2 compendium. The left panel shows the read length (bp), and the right panel displays the library size in millions of reads. Cell lines are colored by their tissue of origin. (**B**) Schematic overview of the computational pipeline used for processing long-read RNA sequencing data. The Nextflow pipeline automates the workflow, starting from raw PacBio reads and proceeding through circular consensus sequencing (pbccs), demultiplexing (lima), refinement to full-length non-chimeric (FLNC) reads (isoseq-refine), alignment (minimap2), sorting (samtools), and finally, transcript discovery and quantification (bambu).

**Figure S2.**
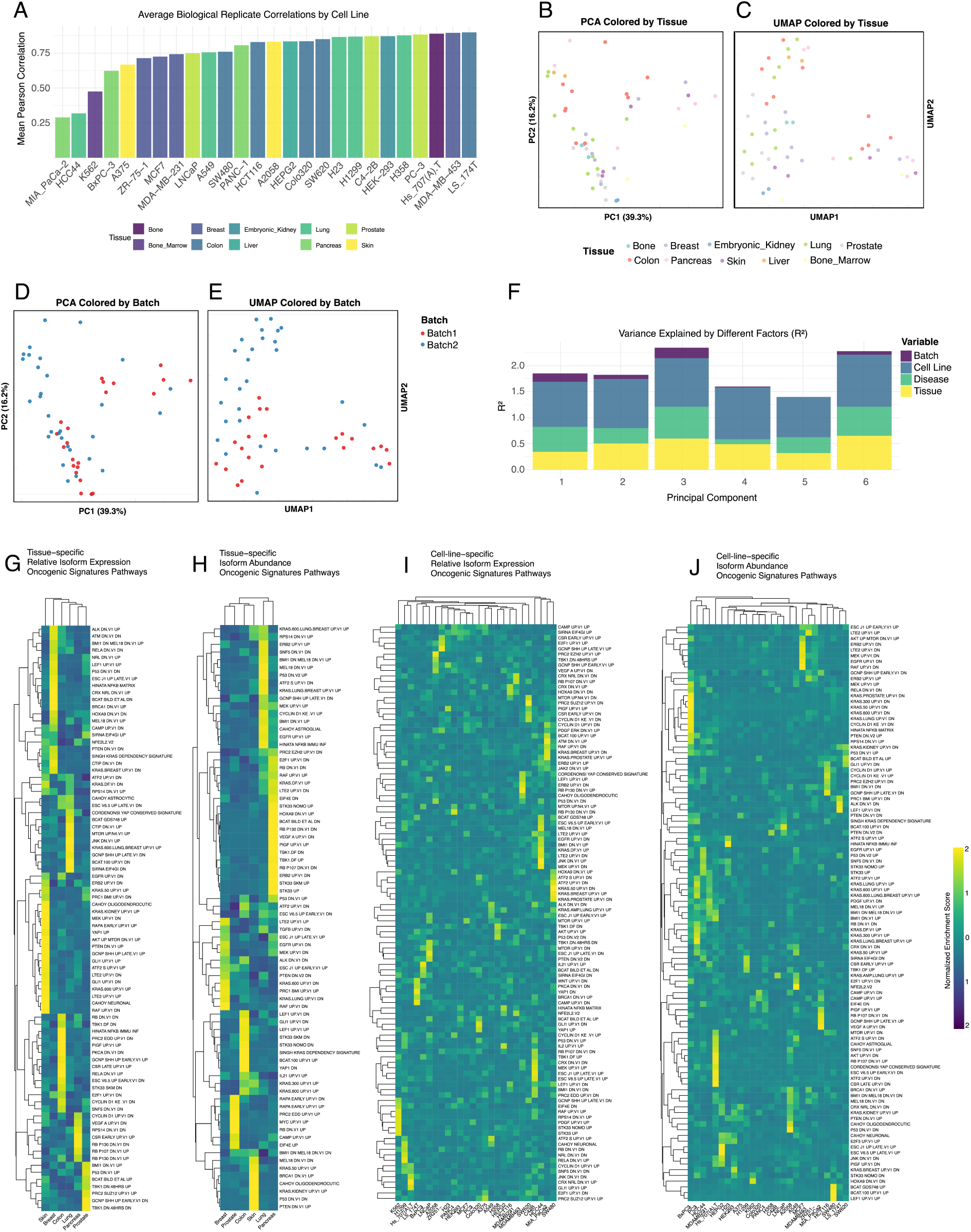
Quality control, exploratory analysis, and functional enrichment analysis of the C2L2 transcriptome compendium. (**A**) Bar plot of average Pearson correlation between biological replicates by cell line (mean = 0.76). (**B**) PCA of the 26 cell lines based on isoform expression, colored by tissue. (**C**) UMAP of the 26 cell lines based on isoform expression, colored by tissue. (**D**) PCA colored by experimental batch. (**E**) UMAP colored by experimental batch. (**F**) Variance decomposition of the top principal components showing *R*_2_ explained by metadata factors, with tissue and cell line as dominant contributors. (**G–J**) Heatmaps of normalized enrichment scores for MSigDB Oncogenic Signature pathways: (**G**) Tissue-specific isoforms (beta regression), (**H**) Tissue-specific isoforms (limma), (**I**) Cell-line-specific isoforms (beta regression), (**J**) Cell-line-specific isoforms (limma).

**Figure S3.**
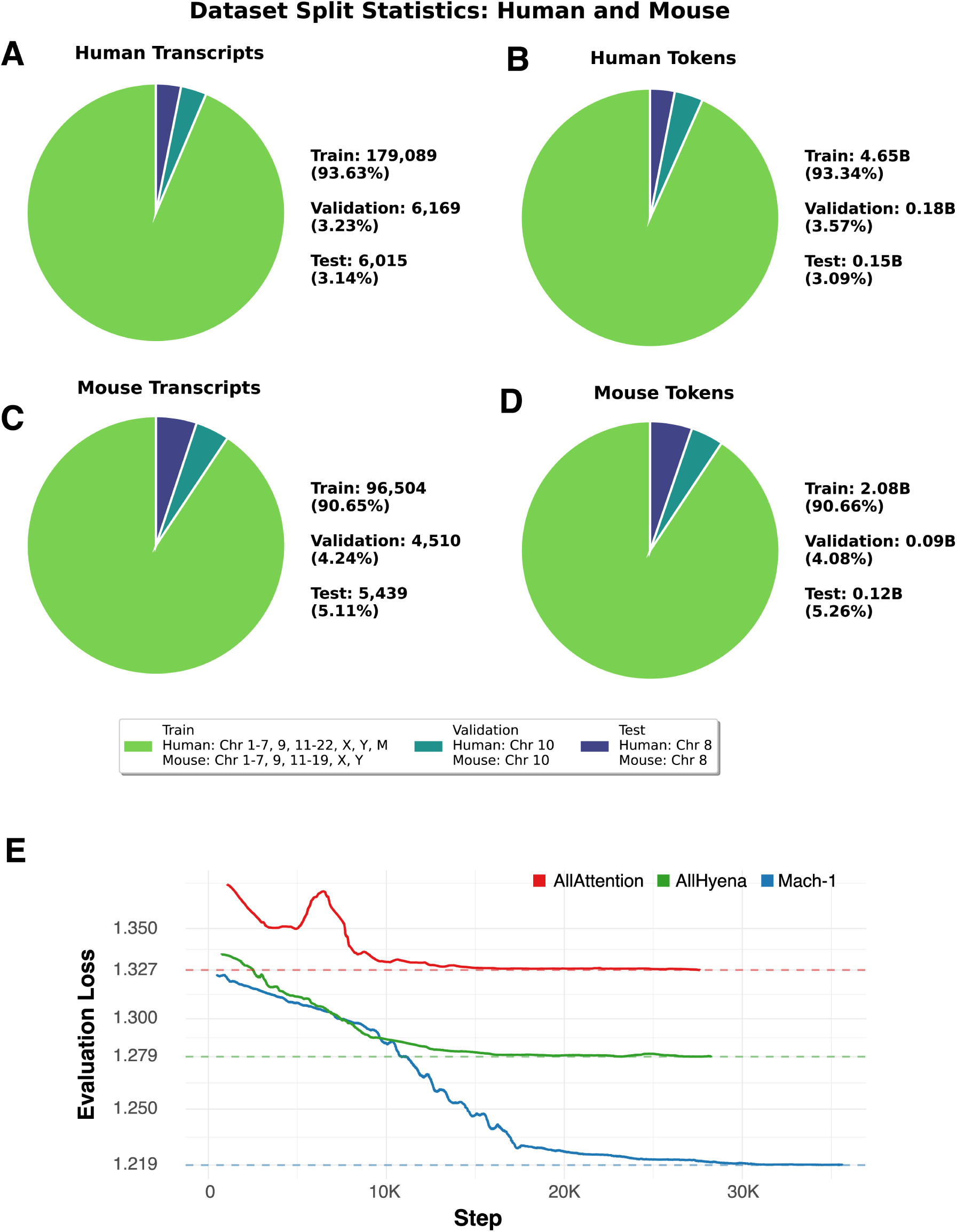
Distribution of transcripts/tokens across splits and model performance. (**A**) Human transcript distribution across splits. (**B**) Human token distribution. (**C**) Mouse transcript distribution. (**D**) Mouse token distribution. (**E**) Evaluation loss curves for Mach-1, All-Hyena, and All-Attention. Mach-1 achieves lowest evaluation loss (1.219).

**Figure S4.**
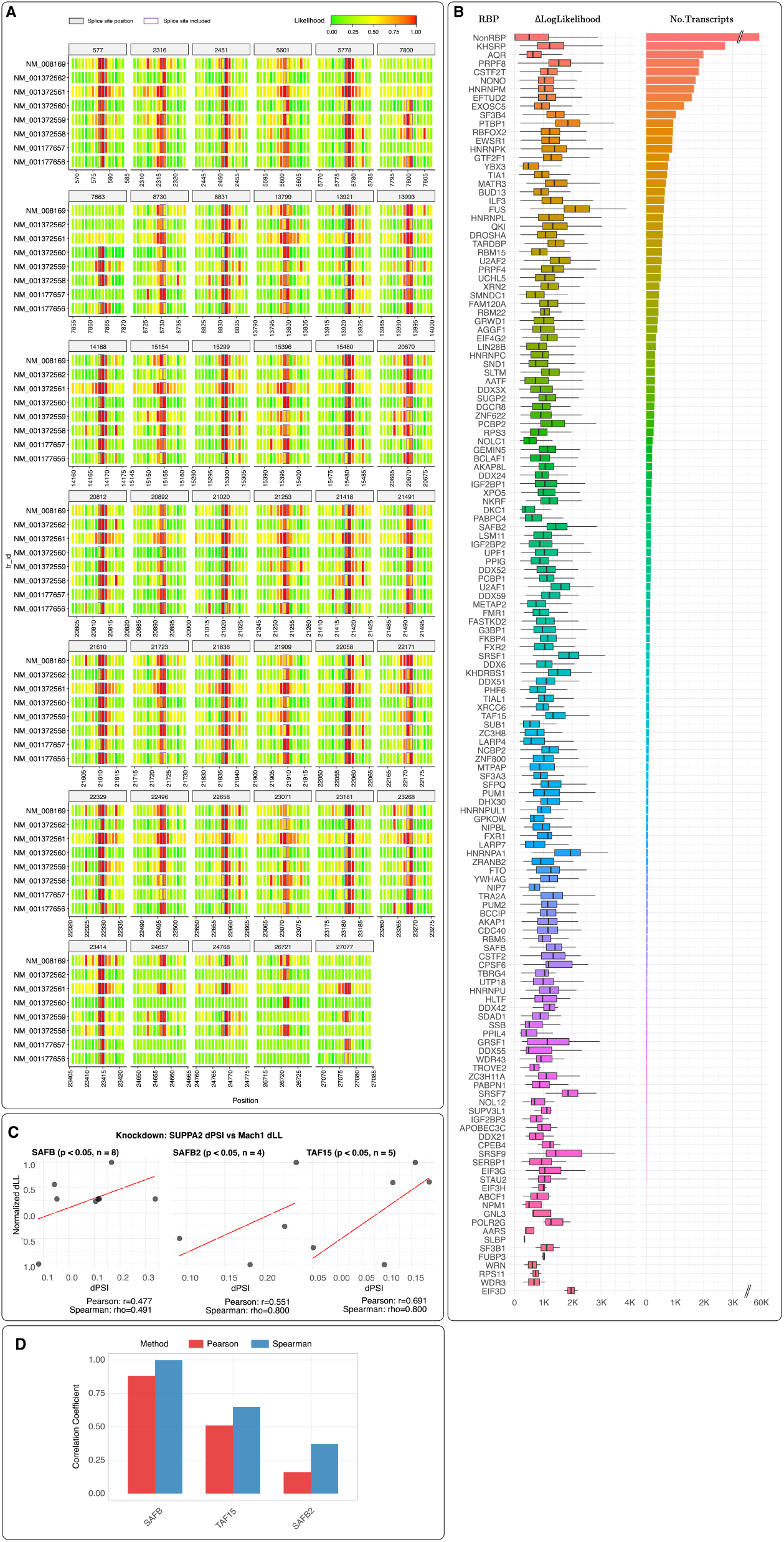
Experimental interrogation of Mach-1’s learned splicing code. (**A**) Likelihood scores across *Grin1* transcripts in mouse. (**B**) Distribution of ΔLL scores for *in silico* RBP binding site mutations. (**C**) Correlation between ΔPSI (measured) and ΔLL (predicted) for SAFB/SAFB2/TAF15 knockdowns. (**D**) Pearson and Spearman correlation between ΔLL and logFC of parent transcripts upon RBP knockdown.

**Figure S5.**
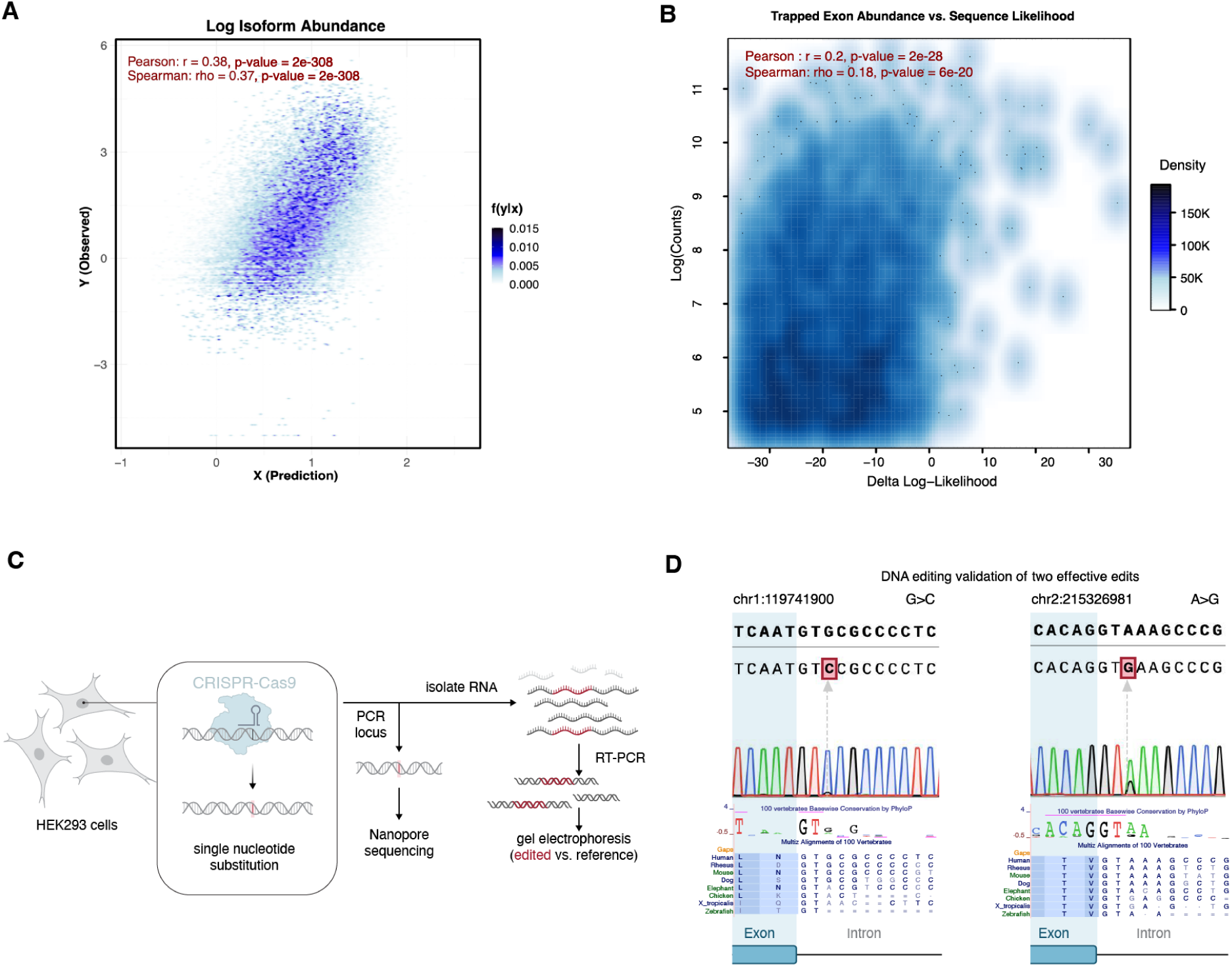
Experimental validation of a VUS via CRISPR-Cas9 genome editing. (**A**) Correlation between predicted and observed expression using PAS embeddings (corr = 0.38). (**B**) Correlation between trapped exon abundance and sequence likelihood (corr = 0.20). (**C**) Schematic of CRISPR-Cas9 editing and validation workflow. (**D**) Nanopore sequencing results confirming two Mach-1-nominated SNP edits, with conservation tracks.

**Figure S6.**
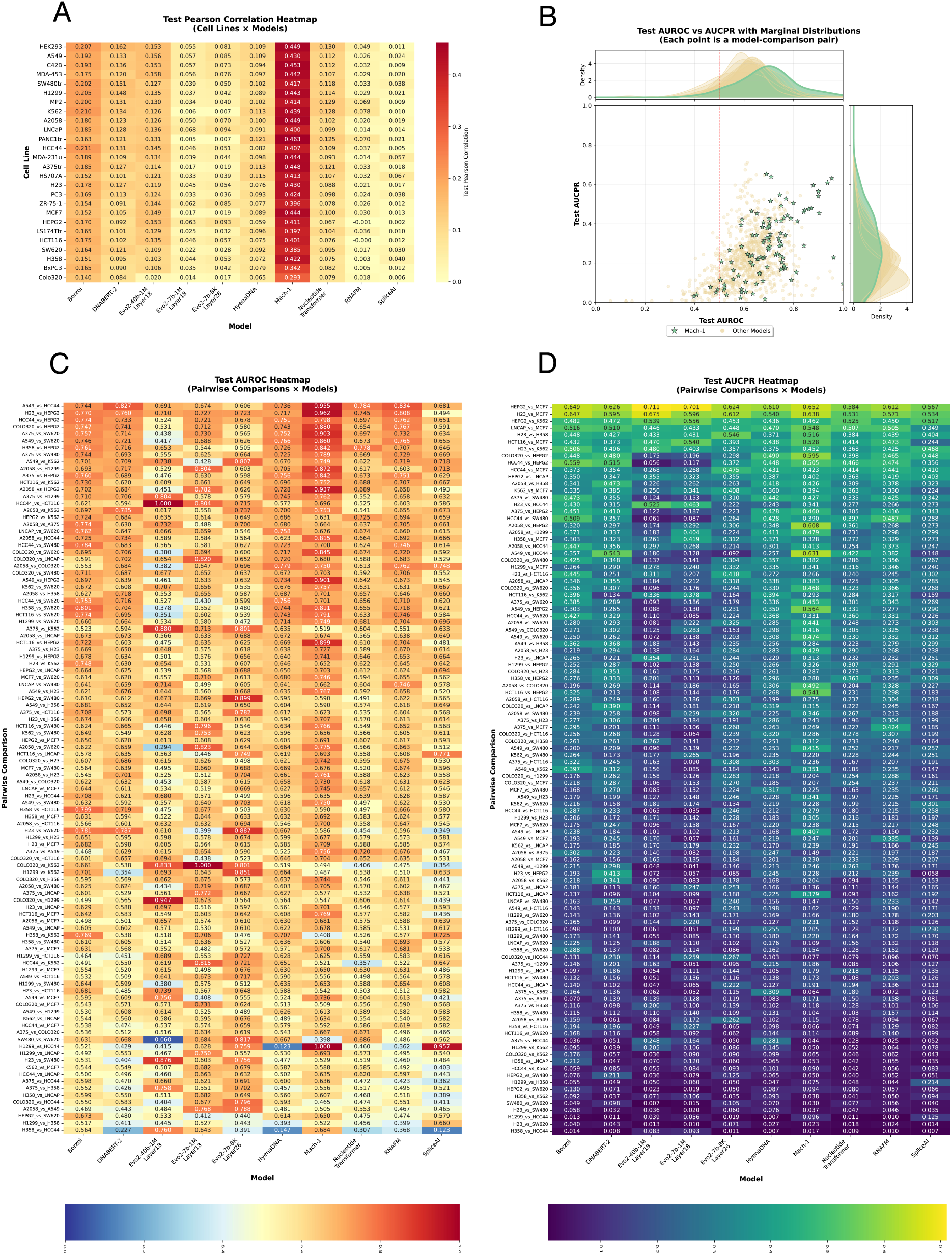
Comprehensive performance of fine-tuned models on cell line-specific prediction tasks. (**A**) Heatmap of Pearson correlations for isoform abundance prediction across cell lines and models. (**B**) Scatter plot comparing Mach-1 with others on pairwise DE task (AUROC, AUPRC). (**C**) Heatmap of AUROC scores across cell-line pairwise DE classification. (**D**) Heatmap of AUPRC scores corresponding to (C).

**Figure S7.**
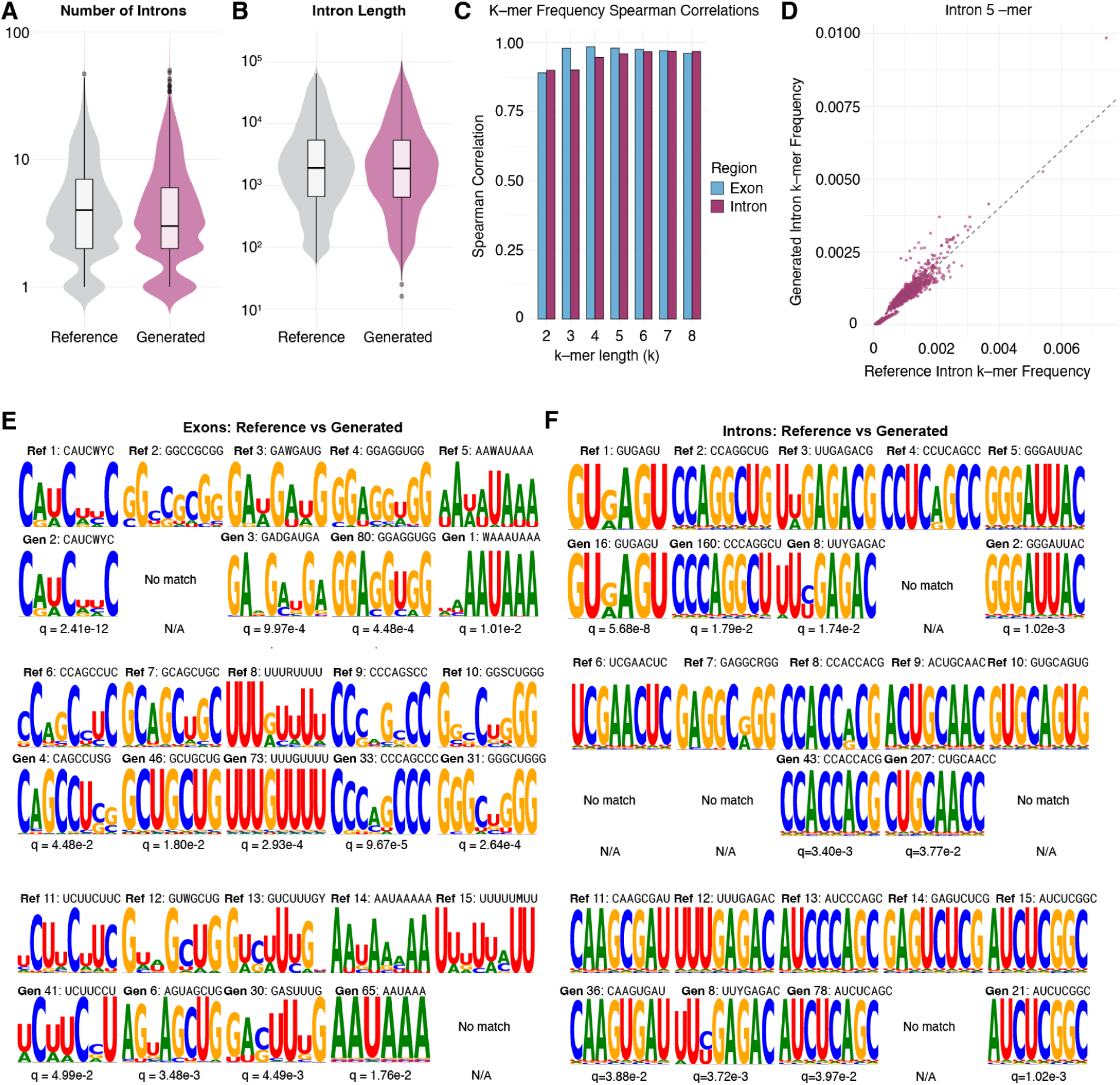
Comparison of generated and reference transcript features. (**A–B**) Violin plots comparing number of introns and intron length between generated and reference sets. (**C**) Bar plot showing Spearman correlation of k-mer frequencies (*k* = 2 to 8) for exonic and intronic regions. (**D**) Scatter plot of 5-mer frequencies in introns (generated vs. reference). (**E–F**) Sequence logos for enriched motifs in exonic and intronic regions, with q-values.

## References

[2310.18780] laughing hyena distillery: Extracting compact recurrences from convolutions, a. URL https://arxiv.org/abs/2310.18780.

Paving the way to efficient architectures: StripedHyena-7B, open source models offering a glimpse into a world beyond transformers, b. URL https://www.together.ai/blog/stripedhyena-7b.

T. Annese, R. Tamma, S. Ruggieri, and D. Ribatti. Angiogenesis in pancreatic cancer: pre-clinical and clinical studies. Cancers, 11(3):381, 2019.

M. Apostolides, B. Choi, A. Navickas, A. Saberi, L. M. Soto, H. Goodarzi, and H. S. Najafabadi. Accurate isoform quantification by joint short-and long-read rna-sequencing. BioRxiv, 2024.

R. Arafeh, T. Shibue, J. M. Dempster, W. C. Hahn, and F. Vazquez. The present and future of the cancer dependency map. Nature Reviews Cancer, 25(1):59–73, 2025.

K. F. Au, V. Sebastiano, P. T. Afshar, J. D. Durruthy, L. Lee, B. A. Williams, H. van Bakel, E. E. Schadt, R. A. Reijo-Pera, J. G. Underwood, and W. H. Wong. Characterization of the human ESC transcriptome by hybrid sequencing. Proceedings of the National Academy of Sciences of the United States of America, 110(50):E4821–30, dec 2013. doi: 10.1073/pnas.1320101110. URL 10.1073/pnas.1320101110.

Ž. Avsec, N. Latysheva, J. Cheng, G. Novati, K. R. Taylor, T. Ward, C. Bycroft, L. Nicolaisen, E. Arvaniti, J. Pan, et al. Alphagenome: advancing regulatory variant effect prediction with a unified dna sequence model. bioRxiv, pages 2025–06, 2025.

G. Benegas, C. Albors, A. J. Aw, C. Ye, and Y. S. Song. Gpn-msa: an alignment-based dna language model for genome-wide variant effect prediction. bioRxiv, pages 2023–10, 2024.

G. Brixi, M. G. Durrant, J. Ku, M. Poli, G. Brockman, D. Chang, G. A. Gonzalez, S. H. King, D. B. Li, A. T. Merchant, et al. Genome modeling and design across all domains of life with evo 2. BioRxiv, pages 2025– 02, 2025.

E. Calvo-Roitberg, C. L. Carroll, G. Kim, V. Sanabria, S. V. Venev, S. T. Mick, J. D. Paquette, M. Uriostegui-Arcos, J. Dekker, A. Fiszbein, and A. A. Pai. mrna initiation and termination are spatially coordinated. Science, 390 (6769):eado8279, 2025. doi: 10.1126/science.ado8279. URL https://www.science.org/doi/abs/10.1126/science.ado8279.

K. Chen, Y. Zhou, M. Ding, Y. Wang, Z. Ren, and Y. Yang. Self-supervised learning on millions of primary RNA sequences from 72 vertebrates improves sequence-based RNA splicing prediction. Briefings in Bioinformatics, 25(3), mar 2024. doi: 10.1093/bib/bbae163. URL 10.1093/bib/bbae163.

Y. Chen, A. Sim, Y. K. Wan, K. Yeo, J. J. X. Lee, M. H. Ling, M. I. Love, and J. Göke. Context-aware transcript quantification from long-read RNA-seq data with bambu. Nature Methods, 20(8):1187–1195, aug 2023. doi: 10.1038/s41592-023-01908-w. URL 10.1038/s41592-023-01908-w.

F. Cribari-Neto and A. Zeileis. Beta regression in R. Journal of Statistical Software, 34(2):1–24, 2010. doi: 10.18637/jss.v034.i02.

H. Dalla-Torre, L. Gonzalez, J. Mendoza-Revilla, N. Lopez Carranza, A. H. Grzywaczewski, F. Oteri, C. Dallago, E. Trop, B. P. de Almeida, H. Sirelkhatim, et al. Nucleotide transformer: building and evaluating robust foundation models for human genomics. Nature Methods, 22(2):287–297, 2025.

A. Fiszbein, K. S. Krick, B. E. Begg, and C. B. Burge. Exon-mediated activation of transcription starts. Cell, 179(7):1551–1565, 2019.

L. T. Gehman, P. Stoilov, J. Maguire, A. Damianov, C.-H. Lin, L. Shiue, M. Ares, I. Mody, and D. L. Black. The splicing regulator rbfox1 (A2BP1) controls neuronal excitation in the mammalian brain. Nature Genetics, 43(7):706–711, may 2011. doi: 10.1038/ng.841. URL 10.1038/ng.841.

L. A. Gilbert, M. A. Horlbeck, B. Adamson, J. E. Villalta, Y. Chen, E. H. Whitehead, C. Guimaraes, B. Panning, H. L. Ploegh, M. C. Bassik, et al. Genome-scale crispr-mediated control of gene repression and activation. Cell, 159(3):647–661, 2014.

S. Gupta, J. A. Stamatoyannopoulos, T. L. Bailey, and W. S. Noble. Quantifying similarity between motifs. Genome Biology, 8(2):R24, 2007. ISSN 1465-6914. doi: 10.1186/gb-2007-8-2-r24. URL 10.1186/gb-2007-8-2-r24.

L. Heinke. A novel g9a–prc2 complex silences developmental genes in prostate cancer. Nature Reviews Molecular Cell Biology, 24(2):84–84, 2023.

C. Huang, R. W. Shuai, P. Baokar, R. Chung, R. Rastogi, P. Kathail, et al., and N. M. Ioannidis. Personal transcriptome variation is poorly explained by current genomic deep learning models. Nature Genetics, 55: 2056–2059, 2023. doi: 10.1038/s41588-023-01574-w. Published 30 November 2023; Open access.

K. Jaganathan, S. Kyriazopoulou Panagiotopoulou, J. F. McRae, S. F. Darbandi, D. Knowles, Y. I. Li, J. A. Kosmicki, J. Arbelaez, W. Cui, G. B. Schwartz, E. D. Chow, E. Kanterakis, H. Gao, A. Kia, S. Batzoglou, S. J. Sanders, and K. K.-H. Farh. Predicting splicing from primary sequence with deep learning. Cell, 176(3): 535–548.e24, jan 2019. ISSN 00928674. doi: 10.1016/j.cell.2018.12.015. URL https://linkinghub.elsevier.com/retrieve/pii/S0092867418316295.

M. J. Landrum, J. M. Lee, G. R. Riley, W. Jang, W. S. Rubinstein, D. M. Church, and D. R. Maglott. ClinVar: public archive of relationships among sequence variation and human phenotype. Nucleic Acids Research, 42 (Database issue):D980–5, jan 2014. doi: 10.1093/nar/gkt1113. URL 10.1093/nar/gkt1113.

H. Li. Minimap2: pairwise alignment for nucleotide sequences. Bioinformatics, 34(18):3094–3100, sep 2018. doi: 10.1093/bioinformatics/bty191. URL 10.1093/bioinformatics/bty191.

H. Li, B. Handsaker, A. Wysoker, T. Fennell, J. Ruan, N. Homer, G. Marth, G. Abecasis, R. Durbin, and . G. P. D. P. Subgroup. The sequence alignment/map format and SAMtools. Bioinformatics, 25(16):2078–2079, aug 2009. doi: 10.1093/bioinformatics/btp352. URL 10.1093/bioinformatics/btp352.

J. Linder, D. Srivastava, H. Yuan, V. Agarwal, and D. R. Kelley. Predicting rna-seq coverage from dna sequence as a unifying model of gene regulation. Nature Genetics, 57(4):949–961, 2025.

P. Machanick and T. L. Bailey. MEME-ChIP: motif analysis of large DNA datasets. Bioinformatics, 27 (12):1696–1697, jun 2011. doi: 10.1093/bioinformatics/btr189. URL 10.1093/bioinformatics/btr189.

Á. Nagy, L. S. Pongor, A. Szabó, M. Santarpia, and B. Győrffy. Kras driven expression signature has prognostic power superior to mutation status in non-small cell lung cancer. International Journal of Cancer, 140(4): 930–937, 2017.

E. Nguyen, M. Poli, M. Faizi, A. Thomas, C. Birch-Sykes, M. Wornow, A. Patel, C. Rabideau, S. Massaroli, Y. Bengio, S. Ermon, S. A. Baccus, and C. Ré. HyenaDNA: Long-range genomic sequence modeling at single nucleotide resolution. *arXiv*, nov 2023. doi: 10.48550/arxiv.2306.15794. URL https://arxiv.org/abs/2306.15794.

E. Nguyen, M. Poli, M. G. Durrant, A. W. Thomas, B. Kang, J. Sullivan, M. Y. Ng, A. Lewis, A. Patel, A. Lou, S. Ermon, S. A. Baccus, T. Hernandez-Boussard, C. Re, P. D. Hsu, and B. L. Hie. Sequence modeling and design from molecular to genome scale with evo. BioRxiv, feb 2024. doi: 10.1101/2024.02.27.582234. URL http://biorxiv.org/lookup/doi/10.1101/2024.02.27.582234.

T. W. Nilsen and B. R. Graveley. Expansion of the eukaryotic proteome by alternative splicing. Nature, 463 (7280):457–463, 2010. doi: 10.1038/nature08909.

F. J. Pardo-Palacios, A. Arzalluz-Luque, L. Kondratova, P. Salguero, J. Mestre-Tomás, R. Amorín, E. Estevan-Morió, T. Liu, A. Nanni, L. McIntyre, E. Tseng, and A. Conesa. SQANTI3: curation of long-read transcriptomes for accurate identification of known and novel isoforms. Nature Methods, 21(5):793–797, may 2024. ISSN 1548-7091. doi: 10.1038/s41592-024-02229-2. URL https://www.nature.com/articles/s41592-024-02229-2.

M. Poli, S. Massaroli, E. Nguyen, D. Y. Fu, T. Dao, S. Baccus, Y. Bengio, S. Ermon, and C. Ré. Hyena hierarchy: Towards larger convolutional language models. *arXiv*, 2023a. doi: 10.48550/arxiv.2302.10866. URL https://arxiv.org/abs/2302.10866.

M. Poli, J. Wang, S. Massaroli, J. Quesnelle, R. Carlow, E. Nguyen, and A. Thomas. StripedHyena: Moving Beyond Transformers with Hybrid Signal Processing Models, 12 2023b. URL https://github.com/togethercomputer/stripedhyena.

D. Ray, H. Kazan, K. B. Cook, M. T. Weirauch, H. S. Najafabadi, X. Li, S. Gueroussov, M. Albu, H. Zheng, A. Yang, H. Na, M. Irimia, L. H. Matzat, R. K. Dale, S. A. Smith, C. A. Yarosh, S. M. Kelly, B. Nabet, D. Mecenas, W. Li, R. S. Laishram, M. Qiao, H. D. Lipshitz, F. Piano, A. H. Corbett, R. P. Carstens, B. J. Frey, R. A. Anderson, K. W. Lynch, L. O. F. Penalva, E. P. Lei, A. G. Fraser, B. J. Blencowe, Q. D. Morris, and T. R. Hughes. A compendium of RNA-binding motifs for decoding gene regulation. Nature, 499(7457):172–177, jul 2013. doi: 10.1038/nature12311. URL 10.1038/nature12311.

F. Reese. Characterizing Transcript Diversity Using Long-Read RNA Sequencing. PhD thesis, University of California, Irvine, 2023.

F. Reese, B. Williams, G. Balderrama-Gutierrez, D. Wyman, M. H. Çelik, E. Rebboah, N. Rezaie, D. Trout, M. Razavi-Mohseni, Y. Jiang, et al. The encode4 long-read rna-seq collection reveals distinct classes of transcript structure diversity. BioRxiv, 2023.

P. Rentzsch, D. Witten, G. M. Cooper, J. Shendure, and M. Kircher. Cadd: predicting the deleteriousness of variants throughout the human genome. Nucleic acids research, 47(D1):D886–D894, 2019.

M. E. Ritchie, B. Phipson, D. Wu, Y. Hu, C. W. Law, W. Shi, and G. K. Smyth. limma powers differential expression analyses for rna-sequencing and microarray studies. Nucleic acids research, 43(7):e47–e47, 2015.

J. Salazar, D. Liang, T. Q. Nguyen, and K. Kirchhoff. Masked language model scoring. 2020. URL https://www.amazon.science/publications/masked-language-model-scoring.

A. Sasse, B. Ng, A. E. Spiro, S. Tasaki, D. A. Bennett, C. Gaiteri, P. L. De Jager, M. Chikina, and S. Mostafavi. Benchmarking of deep neural networks for predicting personal gene expression from dna sequence highlights shortcomings. Nature Genetics, 55:2060–2064, 2023. doi: 10.1038/s41588-023-01524-6. Published 30 November 2023.

T. Shen, Z. Hu, S. Sun, D. Liu, F. Wong, J. Wang, J. Chen, Y. Wang, L. Hong, J. Xiao, et al. Accurate rna 3d structure prediction using a language model-based deep learning approach. Nature Methods, 21(12): 2287–2298, 2024.

U. Singh and E. S. Wurtele. orfipy: a fast and flexible tool for extracting orfs. Bioinformatics, 37(18):3019– 3020, 2021.

N. Stepankiw, A. W. H. Yang, and T. R. Hughes. The human genome contains over a million autonomous exons. Genome Research, 33(11):1865–1878, dec 2023. doi: 10.1101/gr.277792.123. URL 10.1101/gr.277792.123.

J. Su, Y. Lu, S. Pan, B. Wen, and Y. Liu. RoFormer: Enhanced transformer with rotary position embedding. *arXiv*, 2021. doi: 10.48550/arxiv.2104.09864. URL https://arxiv.org/abs/2104.09864.

P. J. Sullivan, J. M. Quinn, W. Wu, M. Pinese, and M. J. Cowley. Splicevardb: A comprehensive database of experimentally validated human splicing variants. The American Journal of Human Genetics, 111(10): 2164–2175, 2024.

Y. Tao, Q. Zhang, H. Wang, X. Yang, and H. Mu. Alternative splicing and related rna binding proteins in human health and disease. Signal Transduction and Targeted Therapy, 9(1):26, 2024. doi: 10.1038/s41392-024-01734-2.

J. L. Trincado, J. C. Entizne, G. Hysenaj, B. Singh, M. Skalic, D. J. Elliott, and E. Eyras. SUPPA2: fast, accurate, and uncertainty-aware differential splicing analysis across multiple conditions. Genome Biology, 19(1):40, mar 2018. doi: 10.1186/s13059-018-1417-1. URL 10.1186/s13059-018-1417-1.

E. L. Van Nostrand, G. A. Pratt, A. A. Shishkin, C. Gelboin-Burkhart, M. Y. Fang, B. Sundararaman, S. M. Blue, T. B. Nguyen, C. Surka, K. Elkins, et al. The encode project: Rna binding proteins. Nature, 530:173–178, 2016. doi: 10.1038/nature19312.

A. Vaswani, N. Shazeer, N. Parmar, J. Uszkoreit, L. Jones, A. N. Gomez, L. Kaiser, and I. Polosukhin. Attention is all you need. *arXiv*, 2017. doi: 10.48550/arxiv.1706.03762. URL https://arxiv.org/abs/1706.03762.

Z. Wang and C. B. Burge. Splicing regulation: from a parts list of regulatory elements to an integrated splicing code. RNA, 14(5):802–813, 2008. doi: 10.1261/rna.876308.

Z. Xie, A. Bailey, M. V. Kuleshov, D. J. Clarke, J. E. Evangelista, S. L. Jenkins, A. Lachmann, M. L. Wojciechowicz, E. Kropiwnicki, K. M. Jagodnik, et al. Gene set knowledge discovery with enrichr. Current protocols, 1(3): e90, 2021.

Y. A. Yoo, M. Roh, A. F. Naseem, B. Lysy, M. M. Desouki, K. Unno, and S. A. Abdulkadir. Bmi1 marks distinct castration-resistant luminal progenitor cells competent for prostate regeneration and tumour initiation. Nature Communications, 7(1):12943, 2016.

T. Zeng and Y. I. Li. Predicting rna splicing from dna sequence using pangolin. Genome biology, 23(1):103, 2022.

Z. Zhou, Y. Ji, W. Li, P. Dutta, R. Davuluri, and H. Liu. Dnabert-2: Efficient foundation model and benchmark for multi-species genome. arXiv preprint arXiv:2306.15006, 2023.

